# Phosphorylation Status of B Beta Subunit Controls PP2A activity in Ethylene-mediated Root Growth Inhibition

**DOI:** 10.1101/2022.05.11.491551

**Authors:** Zhengyao Shao, Bo Zhao, Prashanth Kotla, Jackson G. Burns, Jaclyn Tran, Meiyu Ke, Xu Chen, Karen S. Browning, Hong Qiao

**Author notes:** To whom correspondence should be addressed: Hong Qiao, Department of Molecular Biosciences, University of Texas at Austin 2506 Speedway, NMS 5.324, Austin, TX, 78712. **Author Contributions:** Conceptualization, Z.S., B.Z., and H.Q.; Methodology, Z.S., B.Z., C.X., K.B. and H.Q.; Investigation, Z.S., B.Z., P.K., J.B., J.T., M.K., K.B.; Writing – Original Draft, H.Q.; Writing – Review & Editing, Z.S. K.B., and H.Q.; Visualization, Z.S. and H.Q.; Supervision, H.Q.; Funding Acquisition, H.Q.

## Abstract

The various combinations and regulations of different subunits of phosphatase PP2A holoenzymes underlie their functional complexity and importance. We found that phosphorylation status of Bβ of PP2A acts as a switch to regulate the activity of PP2A. In the absence of ethylene, phosphorylated Bβ leads to an inactivation of PP2A; the substrate EIR1 remains to be phosphorylated, preventing the EIR1 mediated auxin transport in epidermis, leading to normal root growth. Upon the ethylene treatment, the dephosphorylated Bβ mediates the formation of A2-C4-Bβ protein complex to activate PP2A, resulting in the dephosphorylation of EIR1 to promote auxin transport in epidermis of elongation zone, leading to root growth inhibition. Altogether, our research revealed a novel molecular mechanism by which the dephosphorylation of Bβ subunit switches on the PP2A activity to dephosphorylate EIR1 to establish EIR1 mediated auxin transport in epidermis in elongation zone for root growth inhibition in response to ethylene.

**Significance Statement:** Root growth is critical to the establishment of planted seedlings. Ethylene plays an important role in the root growth. Yet, the molecular mechanisms that how ethylene regulates the root growth are largely unexplored. This research sheds light on the molecular mechanisms of the determination of the functional specificity and activity of PP2A in response to stimulus in cell type specific manner and reveals the novel molecular mechanism through which ethylene signaling regulates EIR1 dephosphorylation through Bβ dephosphorylation to active EIR1 mediated auxin transport in epidermis in elongation zone, leading to root growth inhibition.

## Introduction

Ethylene is a gaseous phytohormone that regulates various developmental processes and protects plants from both biotic and abiotic stresses ^1–11^. The ethylene signal is perceived on the endoplasmic reticulum (ER) membrane ^12–15^. ETHYLENE INSENSITIVE 2 (EIN2), a key mediator of ethylene signaling, transduces the ethylene signal from the ER to the nucleus via the dephosphorylation, cleavage, and nuclear translocation of its C-terminal domain (EIN2-C) ^16–19^. In the nucleus, EIN2-C and the transcription factor ETHYLENE INSENSITIVE 3 (EIN3) interact with EIN2 NUCLEAR INTERACTING PROTEIN 1 (ENAP1) to regulate histone acetylation and ethylene-induced transcription ^20–22^. The positive feedback regulation that results from EIN3 dimerization is required for both histone acetylation and gene expression that regulates transcription of ethylene-responsive genes^23^. EIN3 modulates a multitude of downstream transcriptional cascades, including a major feedback regulatory circuitry of the ethylene signaling pathway, and integrates numerous connections between hormone-mediated, growth-response pathways ^24–26^. The positive regulatory role of EIN3 in ethylene signaling is well characterized, and more recently EIN3 was shown to repress gene expression by interacting with a transcription repressor specifically regulated by ethylene ^27^.

Responses to ethylene induce various morphological changes that allow plant adaptation to the ever-changing external environment ^28–35^. Seedlings germinated in the dark in the presence of saturated concentrations of ethylene display a characteristic phenotype known as the triple response, which includes the exaggeration of the curvature of apical hooks and the shortening and thickening of hypocotyls and roots ^36, 37^. How ethylene inhibits root growth has been a longstanding question. It is known that ethylene-mediated inhibition of root cell proliferation at the root apical meristem is mainly achieved by restricting epidermal cell expansion^38, 39^. and that ethylene stimulates auxin biosynthesis and basipetal auxin transport toward the elongation zone, where it activates a local auxin response leading to the inhibition of cell elongation ^40, 41^. The current model is that ethylene stimulates auxin biosynthesis in the root tip and auxin transport in the lateral root cap (LRC) and epidermis by AUX1 and PIN-FORMED 2 (PIN2)^42, 43^. A recent study showed that ethylene stimulates auxin biosynthesis and transport in epidermis which is the key step to mediate root growth inhibition, and this epidermis-specific signaling has an impact on the growth of neighboring cells^39^. However, how ethylene stimulates the auxin transport is unknown. Růzicka et al showed that ethylene promotes the expression of AUX1 and PIN2^43^, but elevation is relatively limited, and therefore unlikely to play a substantial role in regulating auxin transport in the ethylene response. One possibility is that these proteins are activated through ethylene mediated posttranslational regulation.

Protein phosphorylation is one of the most important posttranslational regulations that has been showed to play important functions in ethylene mediated root growth inhibition. In the absence of ethylene, the kinase CONSTITUTIVE TRIPLE RESPONSE 1 (CTR1) phosphorylates EIN2-C, leading to an inactivation of EIN2, resulting in a normal root growth ^44–46^. In the presence of ethylene, EIN2-C is dephosphorylated by an unknown mechanism, resulting in cleavage and nuclear translocation of EIN2-C, leading to root growth inhibition. The phosphorylation of aminocyclopropane-1-carboxylic acid (ACC) synthases by mitogen-activated protein kinase (MAPK) and calcium-dependent protein kinase (CDPK) stabilizes the ACC synthases and hence promotes the synthesis of ACC, the precursor of ethylene that induces the root growth inhibition^47–49^. Furthermore, the phosphorylation of CAP BINDING PROTEIN 20 (CBP20) at Ser245 stimulated by ethylene treatment represses MYB DOMAIN PROTEIN 33 (MYB33) expression through miR319b to inhibit root growth^50^. Most of kinases and phosphatases that are involved in ethylene-mediated regulation are still unidentified, and the molecular mechanisms by which the enzymatic activities of these proteins for particular substrate proteins are fine-tuned are largely unknown.

In this study, we demonstrated that the phosphorylation status of Bβ, the regulatory subunit of the PROTEIN PHOSPHATASE 2A (PP2A), acts as a switch to regulate the activity of the phosphatase in dephosphorylation of the auxin transporter ETHYLENE INSENSITIVE ROOT 1 (EIR1) ^51, 52^, promoting auxin transport in epidermis in elongation zone to inhibit root growth in response to ethylene. We found that the PP2A catalytic subunit C4 can interact with the regulatory subunit Bβ and the scaffolding subunit A2 both *in vitro* and *in vivo* to form a protein complex. In the absence of ethylene, Bβ is phosphorylated, whereas in the presence of ethylene, Bβ is dephosphorylated, leading to the formation of A2-C4-Bβ complex. Furthermore, both our genetics and biochemistry experiments demonstrated that EIR1 is the target of PP2A and that the interaction between EIR1 and the PP2A complex is mediated by A2. In the absence of ethylene, the phosphorylation of Bβ results in inactivation of PP2A, and EIR1 remains phosphorylated, preventing the EIR1 mediated auxin transport to epidermis; as a result, no root growth inhibition occurs. In contrast, in the presence of ethylene, the dephosphorylation of Bβ leads to an activation of PP2A phosphatase to dephosphorylate EIR1, promoting the EIR1 mediated auxin transport in epidermis; as a result, the root growth is inhibited. Taken together, our research reveals the novel molecular mechanism through which ethylene signaling regulates EIR1 phosphorylation through Bβ dephosphorylation to active EIR1 mediated auxin transport in epidermis in elongation zone, leading to root growth inhibition.

## Results

### Ethylene-induced dephosphorylation of Bβ is involved in the regulation of root growth

Protein phosphorylation is one of the most important posttranslational regulations that has been showed to play important functions in ethylene mediated root growth inhibition. In the survey of ethylene regulated phosphoproteomics, we found that in the absence of ethylene, Bβ subunit was phosphorylated at Serine 460; whereas, in the presence of ethylene, the Bβ subunit was completely dephosphorylated (Fig. 1A). The regulatory B subunits of PP2A have been reported to serve as targeting signals for substrate recruitment and phosphatase activity ^53, 54^. We first examined the expression of *Bβ* in its native promoter driven GUS (*proBβ::GUS)* transgenic plants. We found that *Bβ* was expressed across all the plant tissues and its expression was not altered by ethylene (Supplementary Fig. 1A). No obvious ethylene responsive phenotype was detected from the T-DNA knock-out *Bβ* plants (Supplementary Fig. 1B and 1C). To further evaluate whether the Bβ phosphorylation status change plays any roles in the ethylene response, we generated the plants that contained a wild type Bβ (*Bβox*), the plants that contained a phospho-mimic form of Bβ (*Bβ^S460E^ox*) and the plants with a phospho-dead form of Bβ (*Bβ^S460A^ox*). Independent transgenic lines with a comparable protein expression of Bβ, Bβ^S460E^ or Bβ^S460A^ were obtained (Supplementary Fig. 1D) and used for phenotypic analysis. Compared to Col-0, the overexpression of wild type *Bβ* did not display an obvious alteration in ethylene responsive phenotype (Fig. 1B and 1C). However, *Bβ^S460E^ox* plants displayed an ethylene insensitive phenotype. In contrast, *Bβ^S460A^ox* plants displayed a hyper ethylene sensitive phenotype, specifically in roots, even in the absence of ACC (Fig. 1B and 1C). These results suggest that the phosphorylation of Bβ at Ser460 negatively regulates ethylene-mediated root growth inhibition, whereas the dephosphorylated Bβ at Ser460 plays a positive role in the ethylene-mediated root growth inhibition.

**Fig. 1.**
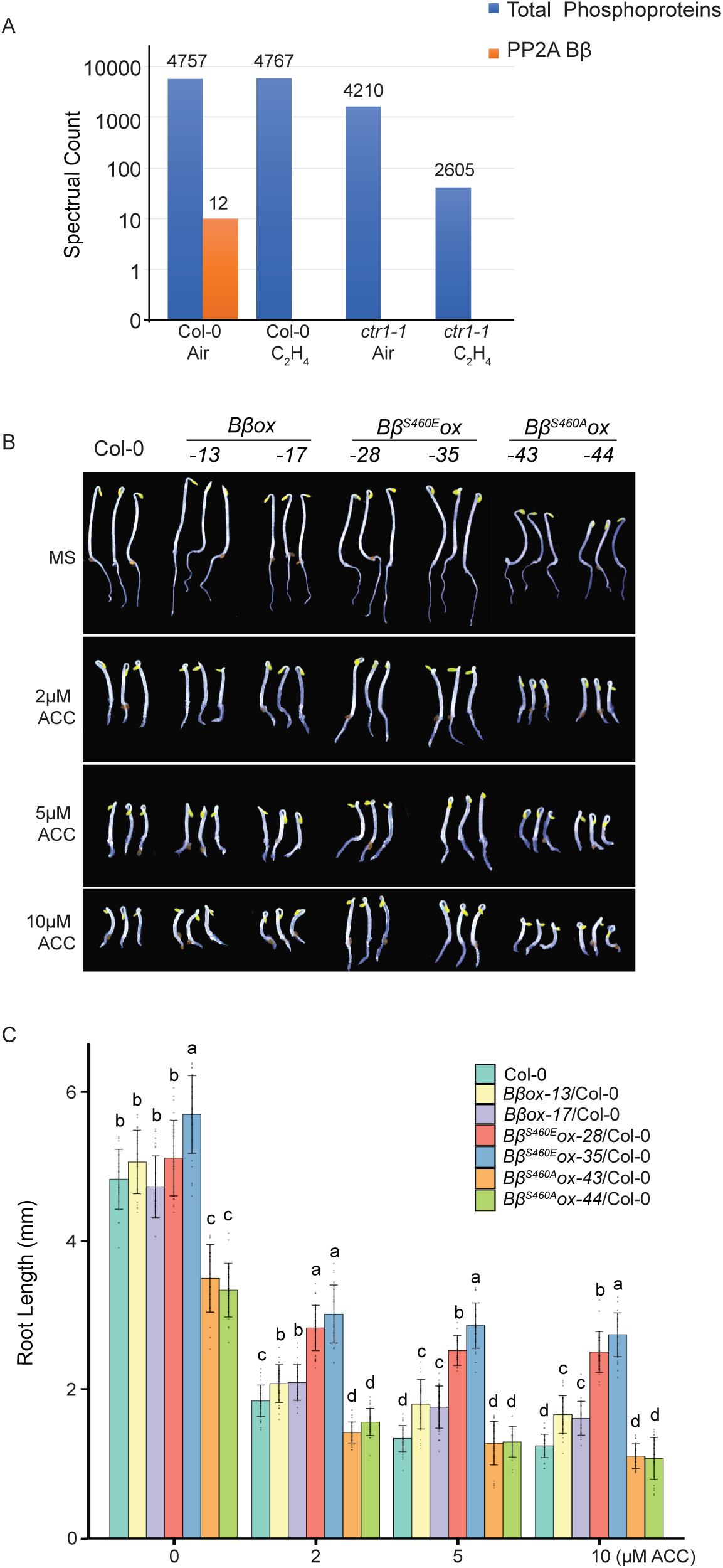
PP2A-Bβ is dephosphorylated at Ser460 upon ethylene treatment, which is involved in the ethylene-mediated root growth. (**A**) Spectral counts of phosphorylated PP2A-Bβ peptides in Col-0 or *ctr1-1* mutants treated with 4 hours of air or ethylene gas. Spectral counts were calculated by averaging three biological replicates. Total spectral counts of all phosphoproteins in each sample serve as an internal control. (**B**) The seedling phenotypes of overexpression of different formats of Bβ subunit. Wild type *B*β, the phospho-mimic *B*β_S460E_, and phospho-dead *B*β*^S460A^* were transformed into Col-0 background respectively. Two independent transgenic lines from each transformation with similar *B*β protein expression levels were selected for phenotypic analysis. The seedlings were grown on MS medium containing 2μM, 5μM or 10μM ACC or without ACC in the continuous dark before being photographed. (**C**) Measurements of the root lengths from the indicated etiolated seedlings described in Fig. 1B. Values are means ± SD of at least 30 seedlings. Different letters indicate significant differences between different genotypes calculated by a two-tailed *t* test with *P* ≤0.05. Individual data points are plotted.

### Bβ is in the same protein complex as PP2A subunits A2, C4, and Bβ phosphorylation status regulates the complex formation

The heterotrimeric Protein Phosphatase 2A (PP2A) is a protein serine/threonine phosphatase composed of a scaffolding subunit (A), a catalytic subunit (C) and a regulatory subunit (B). The various compositions of different PP2A holoenzymes underlie their functional complexity in cell division, morphogenesis, hormone signaling, and stress response ^55–59^. To investigate whether there is a distinct combination of A and C subunits acting collectively with Bβ subunit in ethylene response, we first performed a yeast two hybrid screening by using Bβ as bait protein. We found that Bβ subunit interacted with C4 subunit and C4 interacted with A2 subunit (Supplementary Fig. 2A and 2B). The interaction was further confirmed by the semi-in vivo assay using A2, Bβ and C4 transiently expressed in the tobacco leaves (Fig. 2A and 2B). To examine whether A2 and C4 are associated with Bβ *in planta* spatially, we examined the gene expression profile of *A2*, *C4* using eFP provided by Tair website (Supplementary Fig. 2C and 2E). Both of them expressed across all the tissues, showing that they shared a similar expression pattern with *Bβ* (Supplementary Fig. 2C, 2D, and 2E).

**Fig 2.**
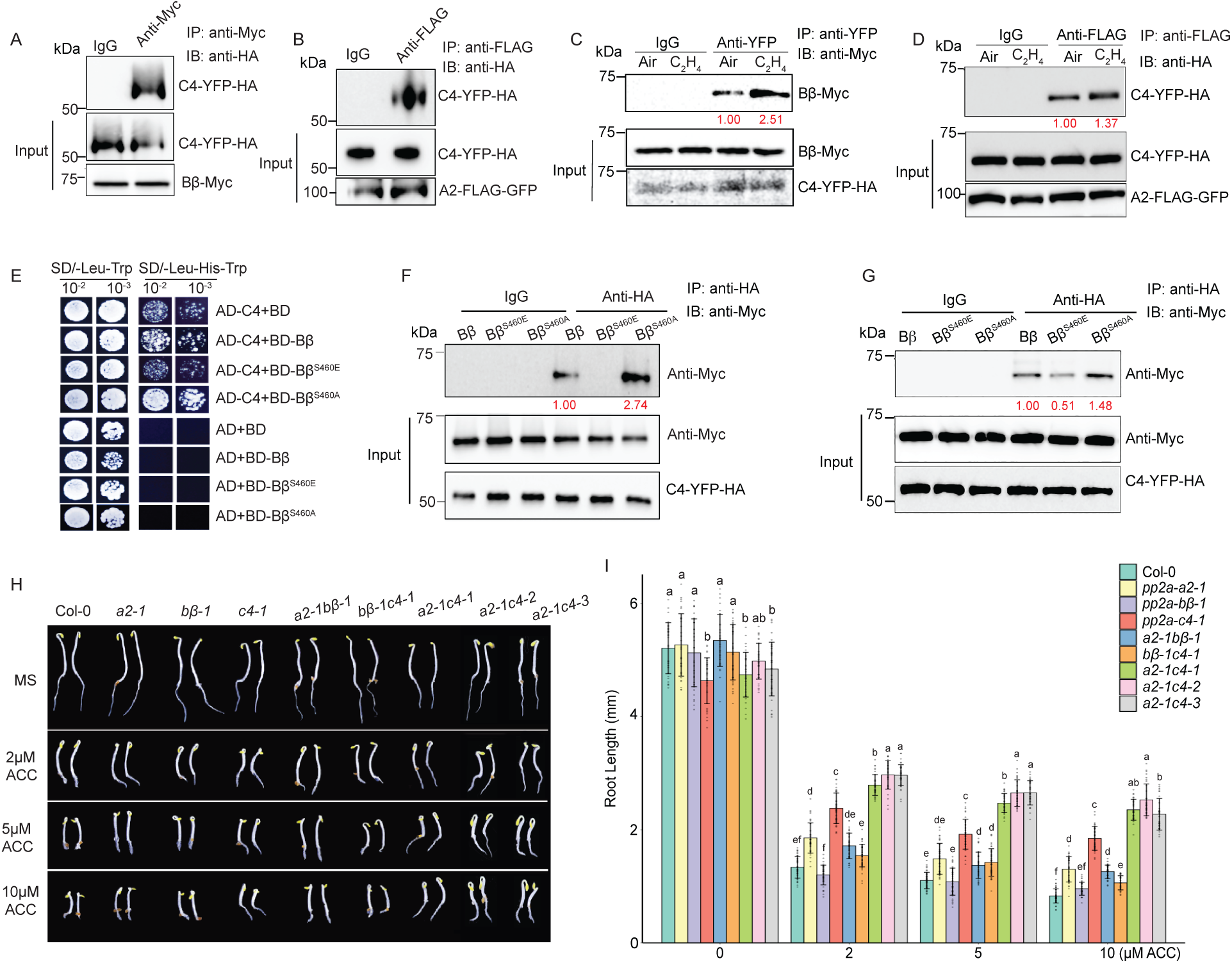
A2,*B*β, C4 subunits are in the same protein complex that involved in ethylene mediated root growth inhibition. (**A** and **B**) Pull-down assays of *B*β (*B*β-Myc) with C4 (C4-YFP-HA) (A) and A2 (A2-FLAG-GFP) with C4 (B) transiently expressed in *N. benthamiana* leaves. (**C** and **D**) *in vivo* co-immunoprecipitation assays of *B*β with C4 and A2 with C4. The total protein extracts from seedlings of crossed *B*β*ox-C4ox* (C) or *A2ox-C4ox* (D) F1 transgenic plants treated with four hours of air or ethylene were immunoprecipitated with anti-HA or anti-FLAG, respectively. The co-immunoprecipitated proteins were detected by western blotting using either anti-Myc or anti-HA. The input serves as the loading control. Red number indicates the quantitation value from anti-Myc or anti-HA divided by the input control. IB, immunoblotting; IP, immunoprecipitation. (**E**) Yeast two-hybrid assay to examine the interaction between C4 with the full length CDS of Bý, the phospho-mimic *B*β^S460E^, and the phospho-dead *B*β^S460A^. BD: GAL4 DNA binding domain; AD: GAL4 activation domain. Left panels: Yeasts grown on two-dropout medium as a control; Right panels: yeast grown on selective three-dropout medium. (**F**) Pull-down assays to examine the interaction between C4 with *B*β, *B*β^S460E^, or *B*β^S460A^. Reactions were performed using total plant extracts from *N. benthamiana* transiently co-expressed C4-YFP-HA with *B*β-Myc, or *B*β^S460E^-Myc, or*B*β^S460A^-Myc respectively. (**G**) *In vivo* co-immunoprecipitation assays to detect the interaction between C4 and different formats of *B*β Reactions were performed using total protein extracts from Col-0 protoplasts transiently co-expressing C4-YFP-HA with *B*β-Myc, *B*β^S460E^-Myc, or *B*β^S460A^-Myc respectively. The total protein extracts were immunoprecipitated with anti-HA antibody, and the anti-Myc antibody was used to detect the indicated proteins in the western blot assay for the indicated proteins. Red numbers indicate the quantitation value from IP product detection in the air or ethylene treatment normalized by their input western blot intensity. (**H-I**) Ethylene responsive phenotypes in the roots of representative plants indicated in the figure. (H)Three-day-old seedlings were grown on MS medium containing 2μM, 5μM and 10μM ACC or without ACC in the dark before being photographed. (**I)** Measurements of root lengths of the plants indicated in (H). Values are means ± SD of at least 30 seedlings. Individual data points of root length measurement are plotted. Different letters represent significant differences between each genotype calculated by a two-tailed *t* test with *P* ≤0.05.

To examine whether and how these subunits interact *in vivo* and to determine whether ethylene influences their interactions, we generated transgenic plants containing both *C4-YFP-HA* and *A2-FLAG-GFP,* or both *C4-YFP-HA and Bβ-Myc.* The *in vivo* co-immunoprecipitation was conducted by using three-day-old etiolated transgenic seedlings treated with or without four hours of ethylene gas. As shown in Fig. 2C and 2D, C4 interacted with both A2 and Bβ, and the interactions were enhanced by the ethylene treatment (Fig. 2C and 2D). Intriguingly, a previous study found that A2, Bβ, and C4 were co-purified by chromatography with a high prominence ^60^, which strongly supports our finding that C4 interacts with both A2 and Bβ to form a PP2A holoenzyme complex.

Since the ethylene treatment enhances Bβ and C4 interaction and Bβ undergoes dephosphorylation in response to ethylene, we aimed to test whether and how the Bβ phosphorylation status influences the interaction between Bβ and C4. In a yeast two hybrid assay, we found that the phospho-dead form of Bβ (Bβ^S460A^) strongly interacted with C4, whereas no interaction was detected between C4 and phospho-mimic form of Bβ (Bβ^S460E^) (Fig. 2E). This result was further supported by a semi *in vivo* co-immunoprecipitation assay of the proteins expressed from tobacco leaves (Fig. 2F). To further validate the interaction *in vivo*, we conducted the co-immunoprecipitation from extracts of protoplasts derived from Col-0 seedlings that were infected with *C4-YFP-HA/Bβ-Myc*, *C4-YFP-HA/Bβ^S460E^-Myc,* or *C4-YFP-HA/Bβ^S460A^-Myc*. Consistent with the *in vitro* assays, the interaction between C4 and Bβ was enhance by the dephosphorylation of Bβ (Fig. 2G), suggesting that the dephosphorylation of Bβ can potentially enhance the formation of A2-Bβ-C4 complex.

To further explore whether A2 and C4 subunits function in the ethylene response genetically, we obtained their T-DNA insertion mutants, *a2-1* and *c4-1,* from the *Arabidopsis* Biological Resource Center ^61, 62^ (Supplementary Fig. 2F and 2G) and examined their ethylene responses in the presence of various concentrations of ACC. We found that the *c4-1* single mutant displayed a moderate ethylene insensitivity in roots (Fig. 2H and 2I). Given the fact that C4 interacts with both Bβ and A2, we then generated *a2-1c4-1* and *bβ-1c4-1* double mutants and examined their ethylene responses. The double mutants displayed different degrees of ethylene irresponsiveness in roots (Fig. 2H and 2I). Notably, the partial ethylene insensitivity observed in *c4-1* roots was significantly enhanced in *a2-1c4-1* double mutant (Fig. 2H and 2I). In order to further confirm the *a2-1c4-1* double mutant phenotype, we generated different alleles of *a2c4* double mutant, named as *a2-1c4-2* and *a2-1c4-3,* by using CRISPR-Cas9 mutagenesis approach to make mutations in *C4* in *a2-1* single mutant background (Supplementary Fig. 2H). Both alleles displayed a similar root phenotype with that of *a2-1c4-1* in response to ethylene, suggesting that A2 and C4 cofunction in the ethylene mediated root growth inhibition.

### A2 and C4 subunits are involved in the ethylene response and EIR1 is the target of A2 subunit

As we screened the potential targets of PP2A by yeast two-hybrid assay, EIR1 appeared to interact with A2 (Supplementary Fig. 3A and 3B). This interaction was further confirmed by the reciprocal *in vitro* pull-down assays using proteins that were transiently expressed in *Arabidopsis* protoplasts derived from Col-0 plants (Fig. 3A and 3B). To validate this interaction *in vivo* and to examine whether the interaction is regulated by the ethylene treatment, we generated *A2-FLAG-GFP* transgenic plants and conducted *in vivo* immunoprecipitation in the membrane fractions from root tissues of three-day-old etiolated seedlings with and without four hours of ethylene treatment and native antibody against EIR1 was used according to the previous publication ^63^. The interaction between EIR1 and A2 was detected in these plants with and without ethylene treatments, and the interaction between A2 and EIR1 was enhanced in the presence of ethylene (Fig. 3C). Furthermore, different EIR1 species were detected in the samples treated with ethylene from that in untreated samples (Fig. 3C). In the absence of ethylene, EIR1 proteins were of higher molecular weight, suggesting that EIR1 is post-translationally modified in response to ethylene.

**Fig3.**
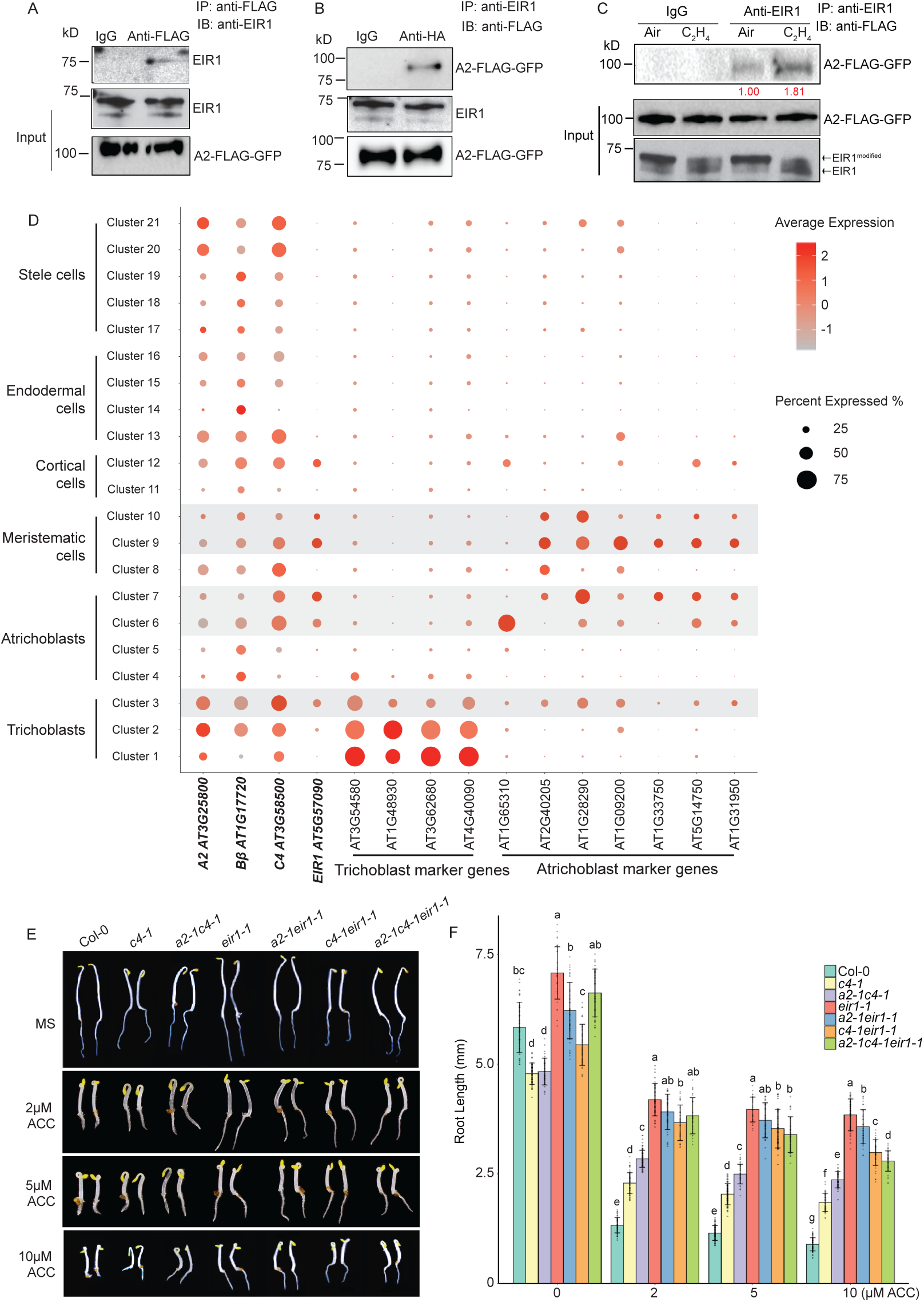
EIR1 is a PP2A target, and A2 and C4 function in ethylene mediated root growth inhibition via EIR1. (**A** and **B**) Reciprocal pull-down assays of A2 (A2-FLAG-GFP) with EIR1 transiently expressed in *Arabidopsis* wild-type protoplasts. The total proteins from the protoplasts infected with *A2-FLAG-GFP* and *EIR1* were applied for the pull-down assays using A2 as bait (A) or EIR1 as bait (B). The pull-down products were detected by western blotting using anti-EIR1 antibody (A) or anti-FLAG antibody. (**C**) *In vivo* co-immunoprecipitation assays of A2 with EIR1. The membrane protein extracts from the roots of three-day-old etiolated transgenic seedlings carrying *35S::A2-FLAG-GFP* treated with four hours of air or ethylene gas were immunoprecipitated with anti-EIR1. The coimmunoprecipitated A2 proteins were detected by western blotting using anti-FLAG antibody. Red numbers indicate the quantitation value from A2 immunoblot band intensity after IP normalized to its input western blot intensity under air or ethylene treatment. **(D)** Spatial expression profiles for A2, *B*β, C4, and EIR1 in sNucRNA-seq dataset. Shadowed clusters (cluster 3, 6, 7, 9, and 10) indicate the shared cell types expressing four genes. Marker genes which have been previously characterized are used to characterize clusters representing trichoblast and atrichoblast. Dot size represents the percentage of cells in which each gene is expressed (% expressed). Dot colors indicate the average scaled expression of each gene in each cell-type cluster with redder colors representing higher expression levels. **(E)** Representative Col-0, *c4-1*, *a2-1c4-1*, *eir1-1*, *a2-1eir1-1*, *c4-1eir1-1*, and *a2-1c4-1eir1-1* mutants were selected for the photograph. Three-day old seedlings were grown on MS medium containing 2μM, 5μM, and 10μM ACC or without ACC before being photographed. (**F**) Measurement of root lengths from the plants indicated in Fig. 3E. Values are means ± SD of at least 30 seedlings. Individual root length data are plotted as dots. Different letters indicate significant differences between different genotypes with *P* ≤0.05 calculated by a two-tailed *t* test.

Given the fact that A2 interacts with EIR1 and mutation of *eir1-1* impairs the ethylene response specifically in roots ^64^, we compared the spatial gene expression patterns of *A2, C4, Bβ* and *EIR1* in different root cell types using single cell sequencing data that are publicly available^65^. We found that *A2, C4, Bβ* were expressed *a*cross different root cell types (Fig. 3D). When we compared their expression levels and cell type specificity in different cell clusters, we found that their expression was significantly overlapped with that of *EIR1* in trichoblasts and atrichoblasts (epidermis), and meristematic cells (Fig. 3D and Supplementary Fig. 3C and 3D), providing an evidence that A2, C4 and EIR1 could co-function in these root cells. To explore the genetic connections between *EIR1* and *A2* or *C4,* we first compared the phenotypes of *eir1-1* and *a2-1c4-1*. We found that *a2-1c4-1* partially phenocopied *eir1-1* in roots in response to ethylene (Fig. 3E and 3F). We then generated *a2-1eir1-1, c4-1eir1-1* and *a2-1c4-1eir1-1* mutants, and examined their ethylene responses in roots. *a2-1eir1-1, c4-1eir1-1*, and *a2-1c4-1eir1-1* mutants had higher levels of ethylene insensitivity than the respective *a2-1, c4-1*, and *a2-1c4-1* mutants, with phenotypes that were similar to that of the *eir1-1* mutant (Fig. 3E and 3F), showing that EIR1 is required for A2 and C4 subunits to regulate the ethylene-mediated root growth inhibition.

### A2, C4, and ethylene induced dephosphorylation of B**β** are required for EIR1 dephosphorylation

Based on the genetics and biochemistry data, we hypothesized that A2 and C4 regulates EIR1 dephosphorylation in the ethylene response. To test this idea, we first examined whether the post translational regulation of EIR1 observed in Fig. 3C is protein phosphorylation by treating EIR1 proteins with calf intestinal alkaline phosphatase (CIP). After CIP treatment, the higher molecular weight EIR1 in Col-0 without ethylene treatment was barely detectable; whereas, the EIR1 band patterns in Col-0 treated with ethylene did not show significant difference from that without CIP treatment (Fig. 4A and 4B). This result indicates that EIR1 is phosphorylated in the absence of ethylene and the levels of phosphorylation are reduced by ethylene treatment. Next, we compared the EIR1 proteins collected from Col-0 and *a2-1c4-1* root tissues treated with or without ethylene. In Col-0, the EIR1 proteins were phosphorylated without the ethylene treatment; the EIR1 proteins were dephosphorylated with the ethylene treatment. In the *c4-1* and *a2-1c4-1* mutants, however, the phosphorylation of EIR1 was not altered by the ethylene treatment; EIR1 was phosphorylated under both conditions (Fig. 4A). A CIP treatment assay further confirmed that the modification of EIR1 detected in the Col-0 without ethylene treatment and in the *a2-1c4-1* mutant under both conditions was phosphorylation (Fig. 4B). Altogether, these data suggest that in wild type plants, EIR1 is phosphorylated in the absence of ethylene, and its dephosphorylation in the presence of ethylene is dependent on A2 and C4.

**Fig4.**
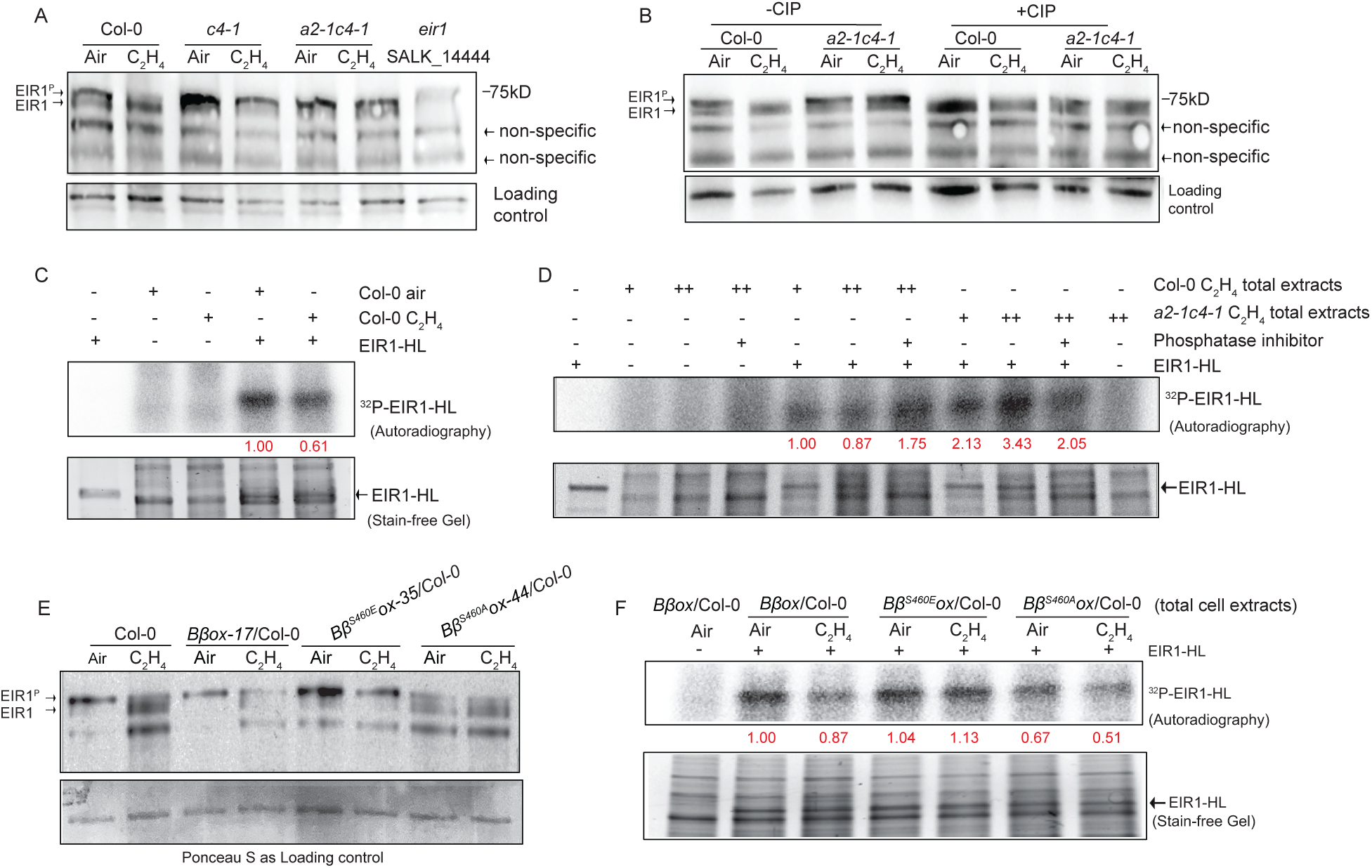
A2,C4 and *B*β^S460E^ are required for the EIR1 dephosphorylation in response to ethylene. (**A** and **B**) Western blot for EIR1 phosphorylation status in membrane fractions from the roots of 3-day etiolated Col-0 and mutant seedlings treated with CIP (B) or not treated with CIP (A) prior to analysis. Blots were probed with anti-EIR1 antibody. Phosphorylated EIR (EIR1^P^) and dephosphorylated EIR1 are labeled. Non-specific bands were used to evaluate loading. (**C** and **D**) Analysis of *in vitro* phosphorylation of EIR1 from total cell lysates from roots of 3-day etiolated Col-0 seedlings treated with either air or ethylene (C) and seedling of the indicated mutants (D). Red numbers indicate the quantitation value from ^32^P-EIR1 phosphor intensity divided by the input intensity shown in the stain-free gel. **(E**) Western blot for EIR1 phosphorylation status in membrane fractions from the roots of 3-day etiolated seedlings indicated in the figure treated with or without ethylene. Blot was probed with anti-EIR1 antibody (upper panel). Ponceau red staining was used as the loading control (bottom panel). (**F**) *In vitro* autoradiograph assay to detect the impact of phosphorylation status of B on the phosphorylation of EIR1 in response to ethylene. *In vitro* purified GST-EIR1-HL was inoculated with total root cell lysates from three-day-old etiolated seedlings from the indicated genotypes with or without 4 hours of ethylene treatment in the presence of MgCl_2_, ATP, and ^32^P-ATP. The reaction samples were subject to SDS-PAGE and the following autoradiography. Red numbers indicate the quantification value from autoradiography of ^32^P-EIR1 phosphor intensity normalized to the input band intensity shown in the stain-free gel. Upper panel: autoradiography; lower panel: stain-free gel as loading control.

Previous studies have shown that the phosphorylation of EIR1 mainly occurs at its central hydrophilic loop (HL) region^66–68^. We therefore examined whether change in EIR1 phosphorylation status in response to ethylene is due to dephosphorylation in this loop. To detect the phosphorylation of EIR1-HL, the purified recombinant GST-EIR1-HL from *in vitro* expression was incubated with the total protein extracted from roots of Col-0 etiolated seedlings treated with air or ethylene and with ^32^P-ATP. Autoradiography showed that phosphorylated ERI1-HL was detected in the samples from plants with and without ethylene treatment (Fig. 4C); however, the EIR1-HL phosphorylation level was drastically reduced in plants treated with ethylene (Fig. 4C). To further verify that the reduction of EIR1 phosphorylation is regulated by A2 and C4 subunits, we examined the phosphorylation of EIR1-HL in total protein extracts from Col-0 seedling and from the *a2-1c4-1* mutant treated with ethylene. The phosphorylation level of EIR1-HL was markedly elevated in the *a2-1c4-1* mutant (Fig. 4D), confirming that ethylene induces the dephosphorylation of EIR1, and this dephosphorylation is A2 and C4 subunits dependent.

Because ethylene mediated Bβ phosphorylation status changing regulates the A2-C4-Bβ complex formation, and phospho-mimicking Bβ^S460E^ and phospho-dead Bβ^S460A^ conferred opposite effects in roots in response to ethylene, we decided to examine whether ethylene-mediated dephosphorylation of Bβ regulates the EIR1 dephosphorylation in response to ethylene. We introduced *Bβ^S460A^ox* and *Bβ^S460E^ox* into *eir1-1* mutant to generate *Bβ^S460A^ox/eir1-1* and *Bβ^S460E^ox/eir1-1*, and examined their ethylene response in roots. We found that the phenotypes conferred by *Bβ^S460A^ox* and *Bβ^S460E^ox* were completely impaired in *Bβ^S460A^ox/eir1-1* and *Bβ^S460E^ox/eir1-1*, and the plants displayed *eir1-1* phenotype (Supplementary Fig. 4A and 4B), showing that EIR1 is required for Bβ to regulate root growth in response to ethylene. We then conducted an immunoblot assay to examine whether Bβ phosphorylation status regulates the phosphorylation of EIR1 protein. In *Bβox* plants, the changes of EIR1 phosphorylation in response to ethylene were similar to that in Col-0 (Fig. 4E). In the *Bβ^S460E^ox* plants, majority of EIR1 proteins were phosphorylated in the absence of ethylene, but the phosphorylation was not altered by the ethylene treatment (Fig. 4E). In contrast, the EIR1 proteins were largely dephosphorylated in the *Bβ^S460A^ox* plants even without the ethylene treatment, and the phosphorylation level was similar to that in the wild-type plants or in the *Bβox* plants that had been treated with ethylene (Fig. 4E). We then conducted *in vitro* EIR1-HL phosphorylation assay by inoculating the *in vitro* purified EIR1-HL with the total protein extracts from the plants of *Bβox, Bβ^S460A^ox* or *Bβ^S460E^ox* treated with or without ethylene. We found that the phosphorylation level of EIR-HL was decreased by the ethylene treatment or by *Bβ^S460A^ox* (Fig. 4F). The ethylene-induced reduction of EIR-HL phosphorylation level was impaired in *Bβ^S460E^ox*. All together, these data demonstrate that the ethylene-induced dephosphorylation of Bβ leads to a dephosphorylation of EIR1.

### Bβ regulates PP2A mediated EIR1 dephosphorylation in a manner that depends on A2 and C4 subunits

The A subunit of PP2A is a scaffolding protein that mediates the formation of PP2A holoenzyme^69^. We speculated that A2 and C4 are required for the regulation of Bβ on the dephosphorylation of EIR1 in response to ethylene. To test this possibility, we first introduced Bβ^S460A^ or Bβ^S460E^ into *a2-1c4-1bβ-1* triple mutant to generate *Bβ^S460A^ox/a2-1c4-1bβ-1* and *Bβ^S460E^ox/a2-1c4-1bβ-1* plants; we then compared their root growth with the roots of *a2-1c4-1, Bβ^S460E^ox*/Col-0, and *Bβ^S460A^ox/*Col-0 in response to ethylene. We found that the roots of *Bβ^S460A^ox*/*a2-1c4-1bβ-1* and of *Bβ^S460E^ox/ a2-1c4-1bβ-1* had phenotypes similar to the roots of *a2-1c4-1bβ-1* mutant (Fig. 5A). The enhanced ethylene-induced root growth inhibition in *Bβ^S460A^ox*/Col-0 and the ethylene insensitivity in root growth in *Bβ^S460E^ox*/Col-0 were impaired in the absence of A2 and C4 (Fig. 5A), providing genetic evidence that A2 and C4 are required for the function of Bβ in response to ethylene. Further, EIR1 dephosphorylation induced by ethylene treatment or by the *Bβ^S460A^ox* was eliminated in the absence of A2 and C4 (Fig. 5B and 5C), supporting the conclusion that A2 and C4 are required for Bβ to regulate PP2A activity on EIR1 dephosphorylation in response to ethylene.

**Fig 5.**
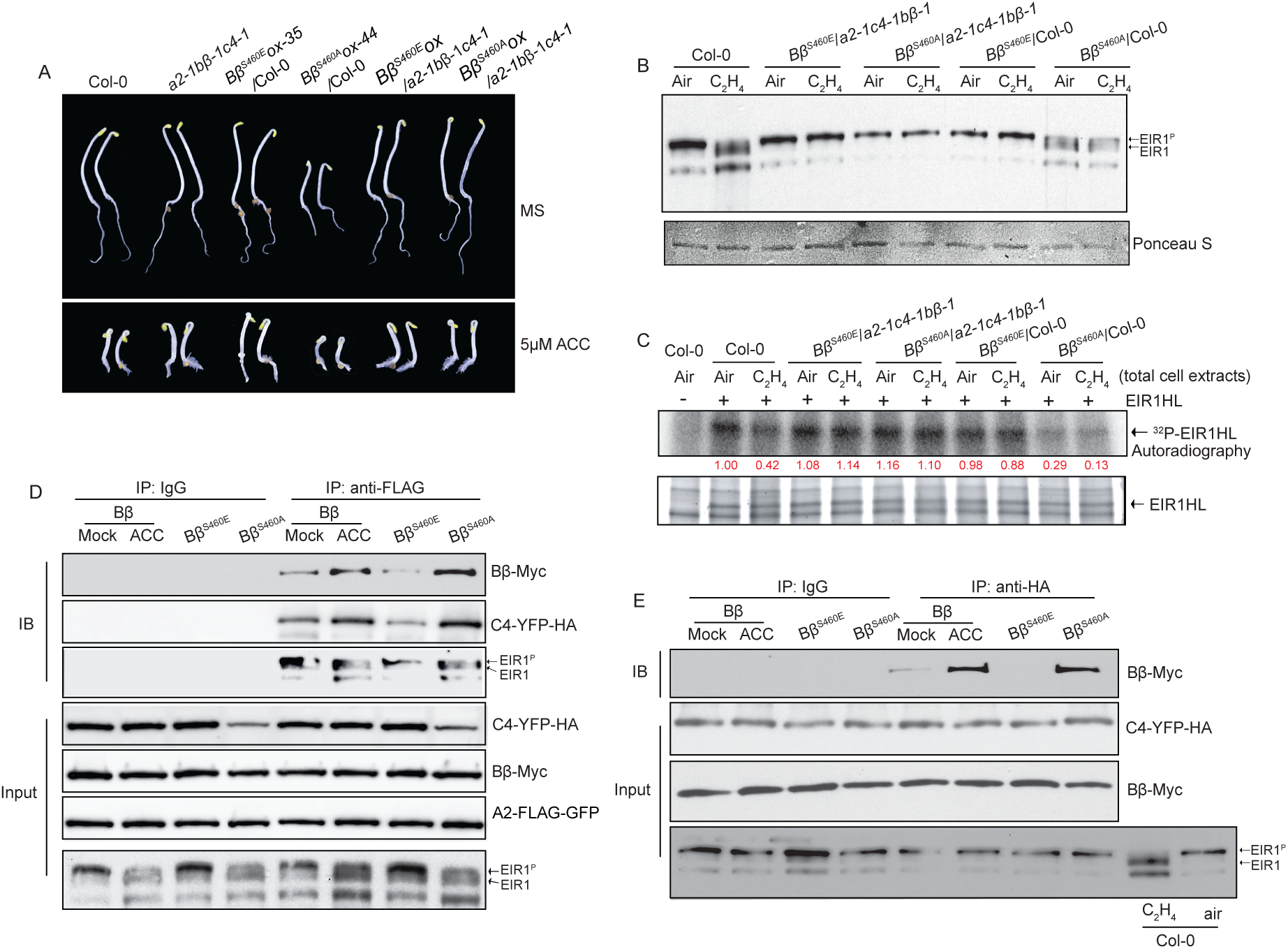
Ethylene dependent dephosphorylation of EIR1 requires A2, C4 and dephosphorylated Bβ. (**A**) The seedling phenotype of the plants indicated in the figure. Three-day-old etiolated transgenic seedlings were grown on MS medium containing 5μM ACC or without ACC in the dark before photographed. (**B**) Western blot analysis of EIR1 phosphorylation status in membrane fractions of Col-0, *B ^S460E^ox*/*a2-1b -1c4-1*, *B ^S460A^ox*/ *a2-1b -1c4-1*, *B ^S460E^ox*/Col-0, and *B ^S460A^ox*/Col-0 three-day-old seedlings treated with or without ethylene. EIR1 was detected with anti-EIR1 antibody (upper panel). Ponceau red staining is used as the loading control (bottom panel). (**C**) *In vitro* autoradiograph assay to examine the function of A2 and C4 in B regulated EIR1 phosphorylation. *In vitro* purified GST-EIR1-HL was inoculated with total root cell lysates from three-day-old etiolated seedlings of indicated genotypes in the presence of purified GST-EIR1-HL, MgCl_2_, ATP, and ^32^P-ATP. The reaction samples were subject to SDS-PAGE and the following autoradiography. Red numbers indicate the quantitation value of individual ^32^P-EIR1 phosphor band intensity normalized to the corresponding input band intensity in each lane shown in the stain-free gel. Upper panel: autoradiography; lower panel: stain-free gel as loading control. (**D**) Western blot analysis of the total cell extracts from the *a2-1b -1c4-1eir1-1* protoplasts that transiently co-expressed A2-FLAG-GFP, C4-YFP-HA, and EIR1 as well as B -Myc, B ^S460E^-Myc, or B ^S460A^-Myc immunoprecipitated with anti-FLAG antibody. (**E**) Western blot analysis of total cell extracts from the *a2-1b -1c4-1eir1-1* protoplasts that transiently co-expressed C4-YFP-HA, and EIR1 as well as B -My, B ^S460E^-Myc, or B ^S460A^-Myc immunoprecipitated with anti-HA antibody.

Based on the data presented above, a model is emerging that A2 mediates the PP2A assembly on the target EIR1 and that the phosphorylation status of Bβ regulates the assembly and the activity of PP2A complex composed of A2, C4, and Bβ, which in turn dephosphorylates EIR1. To test this hypothesis, we conducted co-immunoprecipitation assays to examine whether EIR1 is part of the A2-C4-Bβ complex and whether the phosphorylation status of Bβ will alter the assembly of the complex on EIR1. We first generated the *a2-1bβ-1c4-1eir1-1* quadruple mutant to eradicate the interruptive effects of endogenous A2, Bβ, C4, and EIR1 proteins. We then introduced various combinations of subunits and Bβ mutants into the protoplasts derived from the *a2-1bβ-1c4-1eir1-1* plants: *A2-FLAG-GFP, C4-YFP-HA, EIR1* and *Bβ-Myc*; *A2-FLAG-GFP, C4-YFP-HA, EIR1* and *Bβ^S460A^-Myc;* or *A2-FLAG-GFP, C4-YFP-HA, EIR1* and *Bβ^S460E^-Myc.* The infected protoplasts were then treated with or without four hours of 10µM ACC before testing. All the proteins were well expressed in the *in vitro* assembly system (Fig. 5D). Importantly, the ethylene induced dephosphorylation of EIR1 in the assay was similar to that detected in the Col-0 plants when the four wild-type proteins were expressed (Fig. 5D), showing that the exogenous proteins function properly in the *in vitro* assembly system. We also noticed that the majority of EIR1 was dephosphorylated when the phospho-dead Bβ^S460A^ was expressed, whereas the majority of EIR1 was phosphorylated when the phospho-mimic Bβ^S460E^ was expressed (Fig. 5D). Using A2 as bait, the co-immunoprecipitation assay showed that Bβ, C4, and EIR1 were in the same protein complex (Fig. 5D). In the absence of ACC treatment or when Bβ was constitutively phosphorylated (Bβ^S460E^), the interaction of the protein complex is weakened (Fig. 5D). In contrast, the complex formation was enhanced in the presence of ACC or when Bβ was constitutively dephosphorylated (Bβ^S460A^) (Fig. 5D). We then examined how A2 regulates the A2-Bβ-C4-EIR1 protein complex interaction and the phosphorylation of EIR1 by co-immunoprecipitation in the absence of A2. Under these conditions and using C4 as bait, the interaction between Bβ and C4 was still detectable; however, no EIR1 was detected in the immunoprecipitated products. Furthermore, in the absence of A2, very little dephosphorylated EIR1 was detected from the protoplasts that expressed wild-type Bβ with ACC treatment or in protoplasts that expressed phospho-dead *Bβ^S460A^* (Fig. 5E). These results demonstrate that A2 as a scaffolding protein is required for the assembly and the phosphatase activity of PP2A on EIR1.

### Dephosphorylation of Bβ leads to an activation of EIR1 mediated auxin transport in epidermis in response to ethylene

One way of how EIR1 exclusively restricts root growth is to through EIR1 mediated auxin distribution. Loss-of-function of EIR1 is ethylene insensitive in roots. But our biochemistry and cellular biology data reveal the EIR1 dephosphorylation in ethylene treatment and the EIR1 polarity is not altered by ethylene treatment (Supplementary Fig. 5). Given the dephosphorylation of Bβ leads to a dephosphorylation of EIR1, resulting in root growth inhibition, we speculated that the EIR1 mediated auxin distribution will be regulated by the phosphorylation status of Bβ. To test this idea, we introduced *DR5::GFP*, the proxy of auxin distribution, into *eir1-1, a2-1c4-1, Bβ^S460E^ox and Bβ^S460A^ox,* we then examined the *DR5::GFP* expression in the plants with and without the ethylene treatments and in the plants growing on MS media and MS media supplied with 1 µM ACC. We found that the expression of *DR5::GFP* was limited to the root tip and root cap in the PP2A *a2-1c4-1* double mutant under ethylene treatment or on ACC containing media (Fig. 6A and 6B), and the ethylene induced ectopic expression of *DR5::GFP* in epidermis in Col-0 was impaired in the *a2-1c4-1* mutant (Fig. 6A and 6B), which is similar to that in *eir1-1* mutant (Fig. 6A and 6B). However, in the *Bβ^S460A^ox* plants, where EIR1 is constitutively phosphorylated, the *DR5::GFP* expression was clearly induced in epidermis even in the absence ethylene or ACC. By contrast, the *DR5::GFP* expression was restricted to the root tips and root cap in the *Bβ^S460E^ox* plants, where EIR1 is constitutively dephosphorylated, and the ethylene induces ectopic expression of *DR5::GFP,* proxy of auxin distribution, in epidermis was impaired in the *Bβ^S460E^ox* plants. Fluorescence intensity measurements in epidermal cells in elongation zone in different genetic backgrounds confirmed our observation that ethylene mediated auxin transport in the elongation zone was disrupted *eir1-1, a2-1c4-1,* and *Bβ^S460E^ox* (Fig. 6C, 6D, and 6E). Because the auxin distribution is important for plants gravitropic response, we then did gravity response assay. We found that *a2-1c4-1* and *Bβ^S460E^ox* roots, in which the EIR1 fails to undergo phosphorylation change, had gravitropic defect. The angles of root gravitropic bending for wild-type plants were around 90° after 24 hours gravistimulation, and for *eir1-1* were around 50°. Similar to *etr1-1*, *a2-1c4-1* and *Bβ^S460E^ox* exhibited obvious decreases in bending with the angles ranging from 60° to 70° (Supplementary Fig. 6). These data, together with the result of *DR5::GFP* expression, demonstrate that dephosphorylated EIR1 is critical to proper auxin transport in epidermis in elongation zone in response to ethylene, which requires the activation of PP2A A2-C4-Bβ complex by Bβ dephosphorylation.

**Fig 6.**
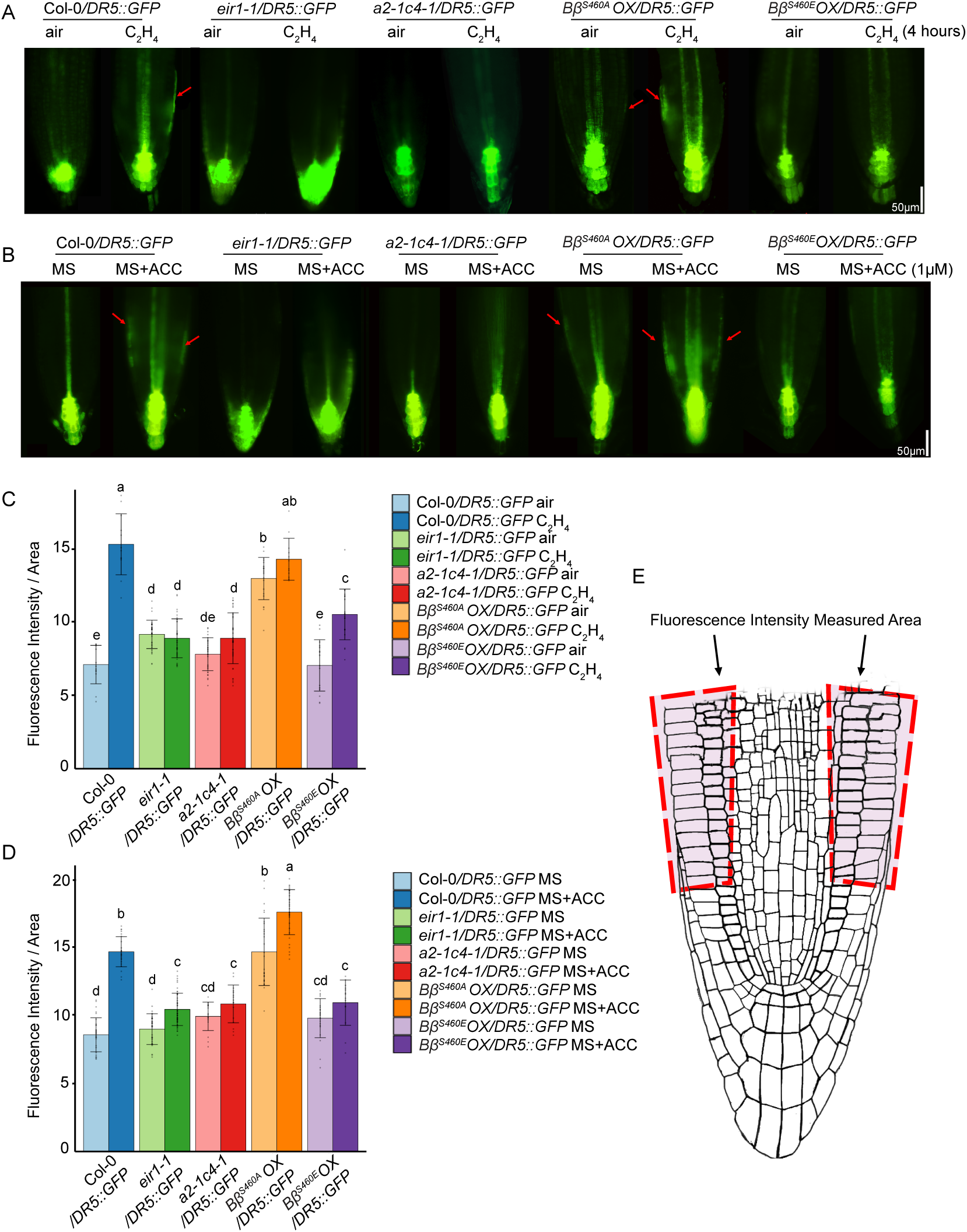
Mutations in PP2A inhibit auxin transport in epidermis in enlogation zone in response to ethylene. **(A and B)** *DR5::GFP* expression in the roots of the plants indicated in the figure with ethylene or ACC treatment. Representative root fluorescence microscope images are displayed. Seedlings were grown on MS medium for 3 days in the dark and photographed after 4 hours of 10ppm ethylene gas or control air treatment (A). Seedings were grown on medium with or without 1µM ACC in the dark for 3 days before photographed (B). The scale bar represents 50μm. (**C** and **D**) Measurement of fluorescence intensity per area in epidermis in the indicated plants under each indicated treatment conditions. Individual measurement data are plotted as dots. Different letters indicate significant differences between different genotypes and treatments with *P* ≤0.05 calculated by a two-tailed *t* test. Values are means ± SD of at least 20 seedlings. (**E**) Root model to indicate root epidermal cells in the elongation zone used for fluorescence measurement.

Put all together, our data support a model that in the absence of ethylene, phosphorylated Bβ at Ser460 destabilizes and subsequentially deactivates the PP2A complex of A2, C4, and Bβ, resulting in EIR1 phosphorylation, preventing auxin transport in epidermis, resulting in normal root growth (Fig. 7). In the presence of ethylene, dephosphorylated Bβ stabilizes A2-C4-Bβ complex, switching on the PP2A activity to dephosphorylate EIR1, activating auxin transport in epidermis, resulting in ethylene mediated root growth inhibition (Fig. 7).

**Fig 7.**
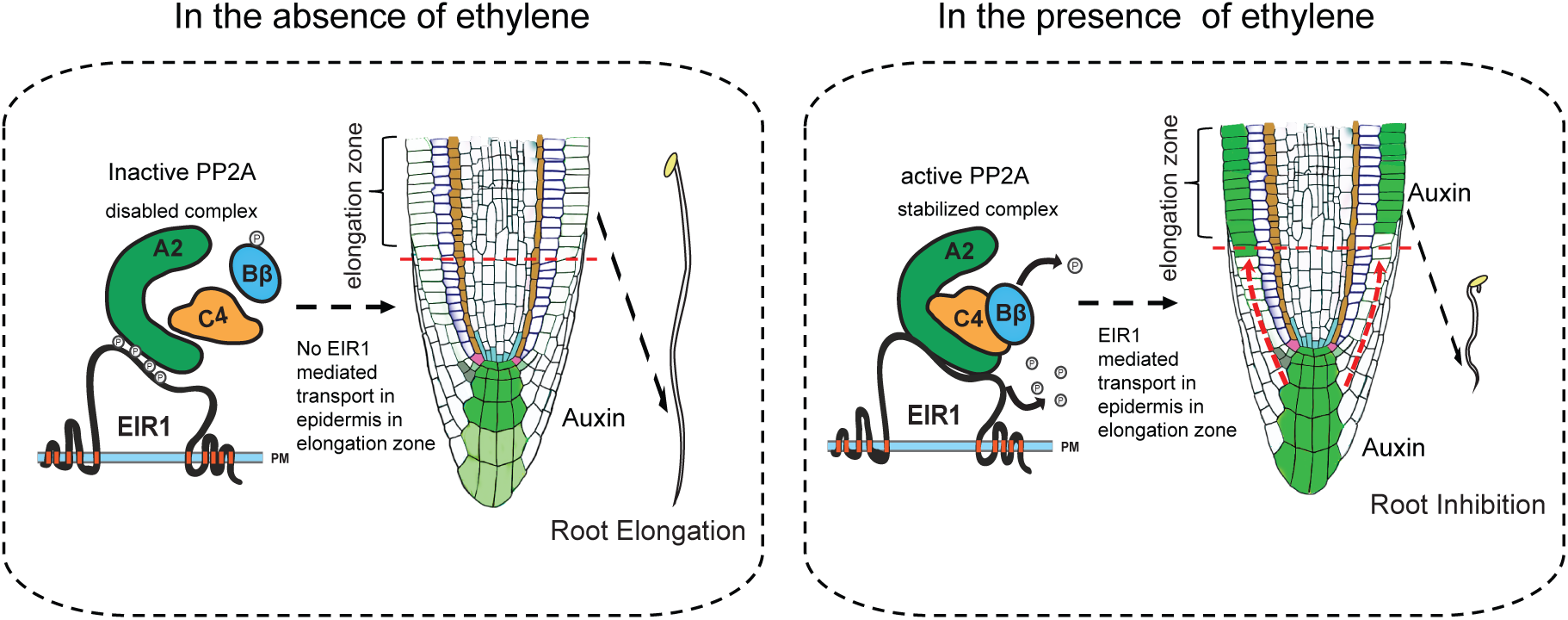
A model to illustrate how ethylene regulates Bβ dephosphorylation to switch on PP2A to dephosphorylate EIR1, leading to root growth inhibition. (Left panel) In the absence of ethylene, phosphorylated B destabilizes and deactivates PP2A complex composed of A2, B, and C4. Phosphorylated EIR1 due to PP2A inactivation prevents auxin transport into epidermis in the root elongation zone, promoting root elongation. (Right panel), In the presence of ethylene, dephosphorylated Bβ stabilizes A2-C4-Bβ complex, switching on the PP2A activity to dephosphorylate EIR1, activating auxin transport in epidermis, resulting in ethylene mediated root growth inhibition.

## Discussion

The PP2A holoenzyme consists of the PP2A core enzyme and a regulatory subunit. The A and C subunits form the PP2A core enzyme, and the regulatory B subunit is the main modulator for PP2A holoenzyme and elicits temporal and spatial specificity. The *Arabidopsis* genome sequence predicts the existence of up to 255 heterotrimeric PP2A isoforms; genes encoding five C subunits, three A subunits, and 17 B subunits have been annotated ^62^. Although the PP2A core enzyme is relatively invariable, the various and interchangeable regulatory B subunits result in a diversity of distinct PP2A holoenzymes. The PP2A has been shown to be involved in numerous biological processes ^53, 57, 59, 60, 70–77^. However, the mechanisms governing the assembly of a distinct PP2A complex to regulate a specific target in a particular biological event are still largely unknown. In this study, we provide multiple lines of compelling evidence that ethylene-mediated dephosphorylation of Bβ switches on the PP2A A2-C4-Bβ activity, which dephosphorylates EIR1, promoting auxin transport in epidermis, leading to ethylene-induced root growth inhibition (Fig. 7). First, our data from biochemistry and genetic experiments revealed that Bβ is dephosphorylated at Ser460 by the ethylene treatment, and Bβ constitutive phosphorylation repressed while Bβ constitutive dephosphorylation enhanced ethylene-mediated root growth inhibition (Fig. 1). Secondly, we showed that PP2A subunit C4 interacts with PP2A A2 and Bβ subunits both *in vitro* and in vivo, and the dephosphorylation of Bβ regulates the formation of A2-C4-Bβ complex (Fig. 2). Thirdly, we demonstrated that EIR1 is one of the targets of PP2A, and A2 mediates the interaction between A2 and EIR1 (Fig. 3). Our genetics data provide a strong piece of evidence that A2, C4 and Bβ function in the same protein complex with EIR1. Biochemistry data showed that ethylene induced dephosphorylation of Bβ leads to activation of PP2A to dephosphorylate EIR1 (Fig. 4 and 5). Finally, we demonstrated that in the absence of ethylene, phosphorylation of Bβ destabilizes the complex of A2, Bβ, and C4 and EIR1 remains phosphorylated, preventing auxin transport in epidermis, leading to normal root growth (Fig. 5 and 6). The dephosphorylation of Bβ in the presence of ethylene switches on the PP2A activity, resulting in dephosphorylation of EIR1, promoting auxin transport in epidermis to inhibit root growth (Fig. 5 and 6).

The B subunit was believed to determine substrate specificity^56, 69, 78^. However, no direct interaction between Bβ and the substrate EIR1 was detected (Supplementary Fig. 3B). Instead, we discovered that the scaffolding subunit A2 directly interacts with EIR1 even in the absence of ethylene (Fig. 3). The dephosphorylation of the regulatory subunit Bβ modulates its interaction with the catalytic subunit C4, leading to an enhanced interaction of these three subunits to assemble the PP2A holoenzyme, and thus activating PP2A function on the dephosphorylation of EIR1 in the presence of ethylene. In this case, it is possible that in the absence of ethylene, phosphorylated Bβ blocks its interaction with C4 to maintain a low level of PP2A activity, therefore preventing the dephosphorylation of EIR1. The interaction between A2 and C4 potentially provides a ready state for PP2A activation. Upon the ethylene treatment, dephosphorylated Bβ can immediately tighten the interaction of the core enzyme A2-C4 and activate its function to dephosphorylate EIR1, which provides an efficient regulation on the specific target in response to the plant hormone ethylene.

It has been proposed that through auxin modulation, ethylene is capable of specifically inhibiting growth of expanding cells or reducing cell proliferation^38, 42^. Vaseva et al., found that the epidermis is the main site of ethylene action controlling plant growth in both roots and shoots^79^. However, the molecular mechanisms are still not clear. Our findings fill one of the gaps between ethylene mediated root growth inhibition and EIR1 mediated auxin transport in epidermis in elongation zone. In the presence of ethylene, Bβ is phosphorylated, switching on PP2A to dephosphorylate EIR1, promoting auxin transport in epidermis in elongation zone, as a result root growth is inhibited. EIR1 was identified as ETHYLENE INSENSITIVE ROOT 1 (also as known as PIN2), whose ethylene insensitivity is restricted in roots ^52^. The phosphorylation of EIR1 is regulated by different plant hormones and nutrient factors ^80–82^. Ötvös et al found that nitrate induced PIN2 dephosphorylation, which affect auxin flux via polarized localization at the plasma membrane ^80^. A very recent study showed that salicylic acid inhibits PP2A activity, leading to a PIN2 hyperphosphorylation, resulting in an increased internalization and a reduced polar membrane localization of PIN2^76^. Particularly, PP2A A1 subunit is responsible for the dephosphorylation of PIN2 in the absence of SA and PIN2 phosphorylation after SA treatment occurs at its hydrophilic loop (HL) region, which is important for the intracellular trafficking of PIN2 ^68^. Unlike salicylic acid, ethylene enhances the activity of PP2A, specifically with the composition of A2-C4-Bβ, to dephosphorylate EIR1. More interestingly, this ethylene regulated phosphorylation also occurs at the hydrophilic loop region (Fig. 4). However, the ethylene induced phosphorylation change of EIR1 does not alter the polarity of EIR1 (Supplementary Fig. 5), which is consistent with a previous publication ^83^, suggesting that ethylene and SA or nitrogen potentially act on different phosphorylation sites of EIR1 to achieve their specific regulations. In case of ethylene treatment, it is possible that the dephosphorylation at specific residues of EIR1 prohibits its biochemical function rather than its polar membrane localization to inhibit auxin transport in epidermis. Identifying the ethylene regulated phosphorylation sites of EIR1 by phosphoproteomic analysis and further analysis the biochemical impacts of ethylene induced dephosphorylation on EIR1 will be the future interests.

## Materials and Methods

### Plant materials, growth condition and hormone responses

*Arabidopsis* (*Arabidopsis thaliana*) accession Columbia (Col-0) was used as the wild type for all the experiments and all of the mutant genotypes in this ecotype background were used. The T-DNA insertion mutants of *pp2a-a2-1* (or *a2-1*, At3G25800, SALK_042724) ^62^, pp2a-*bβ-1* (or *b β*1, At1G17720, SALK_062514), *pp2a-c4-1* (or *c4-1*, At3G58500, SALK_035009) ^61^, *eir1* (AT5G57090, SALK_144447.43.90.x) ^84^, and the point mutation mutant *eir1-1* (CS8085) ^64^ were ordered from Arabidopsis Biological Resource Center. The T-DNA insertions of *a2-1, bβ-1, c4*-1, and eir1 (SALK_144447.43.90.x) were confirmed by genotyping PCR to identify the homozygotes; the *eir1-1* homozygotes were genotyped by Derived Cleaved Amplified Polymorphic Sequences (dCAPS) method ^85^. Primers are listed in Table S1. Seeds were surface sterilized with 50% bleach and 0.01% Triton X-100 for 10 mins, washed with sterile distilled water for four times, sown on Murashige and Skoog medium plates containing 1% sucrose and 1% phytoblend and stratified for three days at 4°C in the dark. For seed propagation, after germination under light, green seedlings were transferred into soil (Promix-HP) and grown in a growth chamber setting with the long-day photoperiod (16 h light/8 h dark) at 22 °C till maturity.

The etiolated seeding triple response assay was performed with sterilized seeds on various concentrations of 1-aminocyclopropane-1-carboxylic acid (ACC, Sigma) plates (0, 2μM, 5μM and 10μM ACC). After three or four weeks upon seed harvesting, approximately 50 seeds of each genotype collected at the similar time were plated on the same MS medium plate per concentration. ACC plates with seeds were placed at 4°C in the dark for three days for stratification and then were exposed to light for four hours and put in the dark for another three days at 22 °C. For phenotypic analysis, representative seedlings from each genotype were selected and placed horizontally on the MS plate and then photographed against a black background. Their root lengths were measured using Fiji ImageJ software ^86^.

For all protein assays, ethylene treatment of *Arabidopsis* etiolated seedlings was performed with sterile seeds growing on MS plates. After stratification and light exposure, MS plates with seeds were placed in air-tight containers in the dark with a flow of hydrocarbon free air at 22°C for three days. Those etiolated seedlings were subsequently treated with ethylene gas at 10 ppm or hydrocarbon-free air for four hours prior to sampling.

### Transient expression in *Nicotiana benthamiana* leaves and in protoplasts

20 mL overnight cultures of *Agrobacterium tumefaciens* (strain GV3101) in LB medium carrying the binary vectors were pelleted by centrifuge at 2000Xg at RT for 15 min and then resuspended in the infiltration buffer (10 mM MgCl_2_, 10 mM MES, pH 5.7, and 100 μM acetosyringone, AS). According to different experiment purpures, pairs of resuspended Agrobacterium cultures were combined and then mixed with an equal volume of p19 resuspension to reach a final OD600 of 0.8 for each resuspension prior to the infiltration at the abaxial side of *N. benthamiana* leaves using a needleless syringe. After growing two or three days in dim light condition, infected N. benthamiana leaves were collected and snap frozen in LN_2_ for further protein analysis.

Four-week-old Arabidopsis plants grown in soil in growth chamber were used for protoplast isolation and the transient expression assay performed according to previous publication ^87^. For protoplast-based transient expressions, protoplasts were transfected with a combination of constructs (approximately 100 μg plasmid DNA in total for 1mL protoplasts for each sample) and incubated at room temperature in the dark for 8 h. ACC was added into the protoplast to reach a final concentration of 10 μM after 4h of incubation. After 4h ACC treatment, protoplasts were harvested by centrifugation at low speed and snap frozen in LN_2_ for further experiments.

### Plasma membrane protein extraction

For EIR1 detection, root tissues from three-day-old etiolated seedlings with air or C_2_H_4_ treatments were harvested and homogenized in LN_2_ to powders. Total membrane extraction was performed as previously^88–90^. In brief, homogenized root materials or pelleted protoplasts were first resuspended in prechilled 1X extraction buffer (50mM Tris-HL, pH= 8.8, 150mM NaCl, 1mM EDTA, 10% (v/v) Glycerol, 1mM PMSF (phenylmethanesulfonylfluoride) and 1X protease inhibitor cocktail) with a combination of phosphatase inhibitors (20mM NaF, 1mM Na_2_MoO_4_, 10mM Na_3_VO_4_, 50mM β-glycerophosphate and 1X PhosStop cocktail (Roche) to preserve EIR1 phosphorylation status. After the addition of equilibrated PVPP (polyvinylpolypyrrolidone) suspension, the total homogenate was centrifuged and cleared at 4°C (2min, 500Xg). Following one time repeat of the extraction with the same volume of extraction buffer, the re-extracted supernatant was combined with the first extract and centrifuged at 4°C for 2h at 22000xg to obtain a total membrane pellet. For EIR1 western blot detection, sample pellets were solubilized with SDS-PAGE Sample Buffer containing 8M urea and 5% SDS and denatured at 50°C for 5 minutes to prevent protein aggregation. For EIR1 related immunoprecipitation (IP), 0.2 mL of membrane solubilization buffer (100mM Tris-HCl, pH = 7.3, 150mM NaCl, 1mM EDTA, 10% glycerol, 20mM NaF, 1% Triton X-100, 1mM Na_2_MoO_4_, 10mM Na_3_VO_4_, 50mM β-glycerophosphate and 1X PhosStop cocktail, 1mM PMSF, 1X protease inhibitor cocktail) was added to the membrane pellet followed by full resuspension. Insoluble particles were removed by centrifugation for two minutes at 22000xg at 4°C. The precleared supernatant containing solubilized proteins was used for the following IP experiments.

### Coimmunoprecipitation and Immunoblot Assays

For co-immunoprecipitation (Co-IP), approximately 2-3 grams of plant material were ground in a prechilled mortar with liquid nitrogen. Soluble total proteins were extracted in two volumes of co-IP buffer (50 mM Tris–Cl, pH 8, 150 mM NaCl, 1 mM EDTA, 0.1% Triton X-100, 1mM PMSF and 1x protease inhibitor cocktail) at 4°C for 15 min with gentle rocking. The lysates were cleared by centrifugation at 4°C (10min, 5000xg). The antibody used for immunoprecipitation was prebound with the equilibrated protein-G coated magnetic beads (Dynabeads, Thermo Fisher) for 3 h with gentle rotation at 4°C. A 10% input aliquot for each co-IP experiment was taken from the cleared supernatant before the rest of the supernatant was added to Dynabeads and incubated at 4°C with gentle rocking overnight. Dynabeads was magnetically precipitated using a DynaMagnetic rack (Thermo Fisher) and then washed five times with 0.5 mL of co-IP buffer. Proteins were then released from Dynabeads using 2× Laemmli sample buffer by heating at 85°C for 8 min.

For protein immunoblots, proteins were separated by SDS–PAGE and transferred to a nitrocellulose membrane (0.2 µm, Bio-rad) by the wet-tank transfer method and blocked with 5% non-fat milk in TBST for 1 h at room temperature before the overnight inoculation in primary antibody at 4°C with gentle rotation. For analysis of EIR1 phosphorylation state, the stacking gel contains 3M urea and the 7% running gel was used and run for 3h at 120V at 4°C. After transfer, the nitrocellulose membranes were blocked in 5% BSA in TBST for 2 h at room temperature prior to immunoblotting. The following antibodies and dilutions were used for immunoblotting: anti-HA (Biolegend, #901503), 1:10000; anti-Myc (CST, #2276), 1:2000; anti-FLAG (rabbit) (CST, #14793), 1:2000; and anti-FLAG (mouse) (Sigma, F3165),1:5000. Goat anti-mouse Kappa Light Chain antibody (Bio-Rad, #105001G) and Goat Anti-Rabbit (IgG (H + L)-HRP Conjugate #1706515, Bio-Rad) were used as secondary antibodies at 1:10000 dilution. Native EIR1 primary antibody was generated by affinity purification of rabbit serum containing -EIR1 antibody kindly provided by Christian Luschnig. HRP activity was detected by enhanced chemiluminescence (ECL, GE Healthcare) according to the manufacturer’s instructions using either a ChemiDoc Imaging System (Bio-Rad) or conventional X-ray films.

### In vitro EIR1-HL phosphorylation assays and CIP treatment

Recombinant GST-EIR1-HL proteins were expressed in the *E. coli* strain BL21 with protein induction when OD600 reached 0.5 with 1 mM IPTG (isopropyl b-D-1-Thiogalactopyranoside) for 4 h at 37°C and then purified over a glutathione Sepharose 4B column (GE Healthcare). The eluted samples were then resolved by SDS-PAGE and stained by Coomassie brilliant blue R-250 to evaluate purity. The in vitro EIR1-HL phosphorylation was performed as previously described ^66^. In brief, crude plant lysates were extracted from homogenized root tissues using 100µL extraction buffer (20 mM Tris-HCl, pH 7.5, 150 mM NaCl, 0.5% Tween-20, 1mM DTT, 1x PMSF, and 1x protease inhibitor cocktail) per sample. Approximately around 1µg of purified GST-EIR1-HL was mixed with kinase buffer (20 mM Tris, pH 7.5, 10mM MnCl_2_, 1 mM ATP and 1mM DTT) containing 1μCi of [γ-^32^P] ATP to a final volume of 10μL. This was added into 10µL crude plant extracts. Reactions were incubated at 30°C for 30min. Reactions were terminated by the addition of 2× SDS loading buffer. After denaturation at 95°C for 2 mins, proteins were resolved on 4-20% stain-free precast gels (Bio-Rad). Either right after the electrophoresis or after Coomassie brilliant blue R250 staining, destaining, and drying, the gels were exposed to phosphor screens and the autoradiograph signal was detected by a Typhoon FLA 9500 system.

For CIP treatment of samples, the membrane pellet from root tissue was first washed five times in extraction buffer without EDTA and phosphatase inhibitors for five times. Then, the pellet was fully resuspended in 50μL CIP buffer (100mM NaCl, 50mM Tris-HCl, pH = 7.9, 10mM MgCl_2_, 1mM DTT) and treated with 2 units of CIP (NEB) at 37°C for 15mins. The reaction was stopped by adding 10μL 6× loading buffer followed by denaturation at 50°C for 5 min. Proteins were resolved on 7% SDS-PAGE gel.

### Quantification and statistical analysis

Details of statistical analyses can be found in the figure legends. R package dplyr was used to perform statistical analyses for phenotypic assays.

### CRISPR-Cas9 mutagenesis

The *a2-1c4-2* and *a2-1c4-3* double mutants were generated by the CRISPR-Cas9 system in *a2-1* single mutant background by following the previous publication^91, 92^. In brief, pairs of two guide RNA (gRNA) sequences targeting *C4* coding region were designed by using CRISPR-PLANT v2 (http://omap.org/crispr2/index.html) online tool. gRNA sequences with high specificity in Arabidopsis genome and predicted efficiency were chosen for PCR amplification using pDT1T2 vector as template and later incorporated into the binary vector pHEE401E. Agrobacterium mediated floral dipping method was used to transform the pHEE401E vector containing C4 gRNA into *a2-1* plants. Genomic DNAs extracted from hygromycin positive T1 plants were used for genotyping and followed by Sanger sequencing to detect the CRISPR-Cas9 directed mutation and obtain double homozygotes of *a2-1c4-2* and *a2-1c4-3*.

### GFP Fluorescence Quantification

The quantification of *DR5*::GFP fluorescence in ethylene and ACC treatment was performed in ImageJ FIJI software (https://imagej.net/software/fiji/). Boxes were drawn surrounding the epidermis in the root elongation zone as area of interest illustrated in Fig. 6E, symmetrically against root axis. Integrated fluorescence intensity and area value were measured for each individual plants and calculated to get mean value for plotting. The significance of differences between different genetic backgrounds and treatment groups were calculated by student *t* test.

### Gravitropic response assay

5-day-old seedlings grown on MS media supplied with 1% sucrose under light day light cycle were used. Plants were transferred to a new vertical plate and aligned prior to the gravistimulation. Pictures were taken 24 hours after the gravitropism assays and the root bending angles were quantified using ImageJ FIJI software with the angle tool function.

### Single nucleus RNA sequencing (sNucRNA-seq) data processing

sNucRNA-seq of Arabidopsis root tissue was obtained from previous publication under accession number GSE155304. Integrated sNucRNA-seq rds file (GSE155304_rnaseq_integration.rds.gz) was used for further analysis of PP2A and EIR1 expression profile. Dot plot and UMAP were generated by the Seurat R package (version 2.3.4) with adaption. Cluster-specific marker genes of trichoblast and atrichoblast cell types were selected based on previous publication^65, 93^.

## Acknowledgments

We thank Dr. C. Luschnig for sharing native anti-EIR1 antibody and C. Jo and A. Basnet for plant and laboratory maintenance. We than Dr D. Kelley for sharing *DR5::GFP* seeds. We thank Dr. J. Xu for sharing *EIR1-GFP* transgenic plants. We thank the Arabidopsis Biological Resource Center for providing seeds. **Funding:** Z.S. was supported by the Graduate Continuing Fellowship from The University of Texas at Austin. This work was supported by grants from the National Institute of Health to H.Q. (NIH-2R01 GM115879-01). **Author contributions:** Conceptualization, Z.S., B.Z., and H.Q.; Methodology, Z.S., B.Z., C.X., K.B. and H.Q.; Investigation, Z.S., B.Z., P.K., J.B., J.T., M.K., K.B.; Writing – Original Draft, H.Q.; Writing – Review & Editing, Z.S. K.B., and H.Q.; Visualization, Z.S. and H.Q.; Supervision, H.Q.; Funding Acquisition, H.Q. **Competing interests:** The authors declare no competing interests. **Data and materials availability:** Further information and requests for reagents and all unique materials generated in this study may be directed to and will be fulfilled by the corresponding author Hong Qiao.

