## supplemental materials for "Phosphorylation Status of B Beta Subunit Controls PP2A activity in Ethylene-mediated Root Growth Inhibition"

Supplemental Figures and Table:

**Supplemental Table 1. A list of primers**

| Name | Forward | Reverse |
|---|---|---|
| *pp2a-a2*_genotyping | CGATGTTACGTGCCCTCTTAC | TCTACCGAATGACCATTTTGC |
| *pp2a-bβ*_genotyping | TTCCATCTCAGGAAACAATGC | AGAAAGAAGCTCTCACCCTGG |
| *pp2a-c4*_genotyping | GCTTGAAAGAACAGCATTTCG | GTGGATTATCACCATCCATCG |
| SALK_144447.43.90.x_genotyping | AAAAACCTAAGAGTTTTGGAAGTG | TTTGTTCAAATTAACGGACCG |
| LBb1.3 | ATTTTGCCGATTTCGGAAC | |
| *eir1-1*_dCAPS_genotyping-1 | TTGCTTGATGTTGTTGATCATTTTATGGGACC | AATCTTTGGTTGCAATGCCATAAATAGACC |
| *eir1-1*_dCAPS_genotyping-2 | TTGCTTGATGTTGTTGATCATTTTATGGGACT | AATCTTTGGTTGCAATGCCATAAATAGACC |
| A2-AD | ACGCGTCGACTATGTCTATGATCGATGAGCC | CTAGACTAGTTTAGCTAGACATCATCACATTG |
| A2-BD | ACGCGTCGACTATGTCTATGATCGATGAGCC | CTAGACTAGTTTAGCTAGACATCATCACATTG |
| Bβ-AD | ACGCGTCGACAATGAACGGTGGTGACG | TCCCCCCGGGTCATGCATAGTACATGTACAAGCT |
| Bβ-BD | ACGCGTCGACAATGAACGGTGGTGACG | TCCCCCCGGGTCATGCATAGTACATGTACAAGCT |
| C4-AD | TCCCCCCGGGAATGGGCGCGAATT | CTAGACTAGTTCAAAGGAAATAGTCAGGTGT |
| C4-BD | TCCCCCCGGGAATGGGCGCGAATT | CTAGACTAGTTCAAAGGAAATAGTCAGGTGT |
| Bβ-pENTR | CACCATGAACGGTGGTGACG | TGCATAGTACATGTACAAGCTGTT |
| C4-pENTR | CACCATGGGCGCGAATT | AAGGAAATAGTCAGGTGTCCT |
| A2-pCambia1300 | CGGGGTACCATGTCTATGATCGATGAGCC | ACGCGTCGACGCTAGACATCATCACATTGTCA |
| EIR1-pCambia1300 | ACGCGTCGACATGATCACCGGCAAAGAC | ACGCGAGCTCTTAAAGCCCCAAAAGAACGTAG |
| EIR1-AD | ACGCGTCGACCATGATCACCGGCAAAGA | CTAGACTAGTAAGCCCCAAAAGAACGTAG |

| | | |
|---|---|---|
| EIR1-BD | ACGCGTCGACCATGATCACCGGCAAAGA | CTAGACTAGTAAGCCCCAAAAGAACGTAG |
| proBβ-GUS-pBI121 | CCCAAGCTTAACTTTTAAACGAGGAGGC | TCCCCCCGGGTTTTTTTTTTTGGCTTTTGGGTTAGATAA |
| Bβ-S460E-mutagenesis | GGATCAGAGGAGCCCGGAACAGAGGCA | TCTGATCCTCGTCTAACCACG |
| Bβ-S460A-mutagenesis | GGATCAGAGGCGCCCGGAACAGAGGCA | TCTGATCCTCGTCTAACCACG |
| C4 CRISPR-Cas9 construct-BsF | ATATATGGTCTCGATTGTTCCACAGAATAATATCCAGTT | |
| C4 CRISPR-Cas9 construct-F0 | TGTTCCACAGAATAATATCCAGTTTTAGAGCTAGAAATAGC | |
| C4 CRISPR-Cas9 construct-R0 | AACGCTGTTGGTTGGCTTGAAACAATCTCTTAGTCGACTCTAC | |
| C4 CRISPR-Cas9 construct-BsR | ATTATTGGTCTCGAAACGCTGTTGGTTGGCTTGAAACAA | |
| C4 CRISPR-Cas9 genotyping | TAATTTTTGGTGTCCATCTACCA | CGACTCAATAGCGTATACTTCAGA |

### Supplemental figures and figure legends

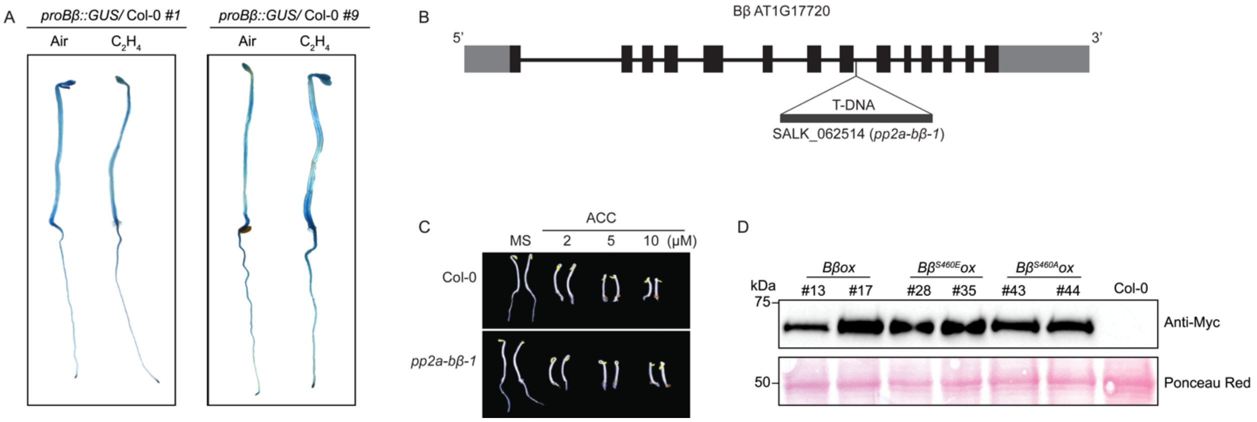


**Figure S1. Related to Figures 1. Bβ expression pattern, PP2A *Bβ* T-DNA insertion mutant, and protein expression levels of *Bβox*/Col-0, *Bβ$^{S460E}$ox*/Col-0 or *Bβ$^{S460A}$ox*/Col-0**

(A) β-glucuronidase (GUS) staining to show the Bβ expression of in Arabidopsis. 3-day-old etiolated independent Col-0 plants carrying *proBβ*::GUS were tested with or without the 4-hours of ethylene treatment. (B) Schematic diagram to show *Bβ* gene (AT1G17720) and the location of the T-DNA insertion in SALK_062514 (*pp2a-bβ-1*). (C) Ethylene response of *bβ-1* mutant. 3-day-old seedlings were grown on MS medium containing 2μM, 5μM and 10μM ACC or without ACC in the dark before being photographed. (D) Western blot assay to show the protein expression in different transgenic plants. The total proteins from the indicated transgenic plants were subjected to the western blot assay with anti-Myc antibody. The ponceau staining serves a loading control.

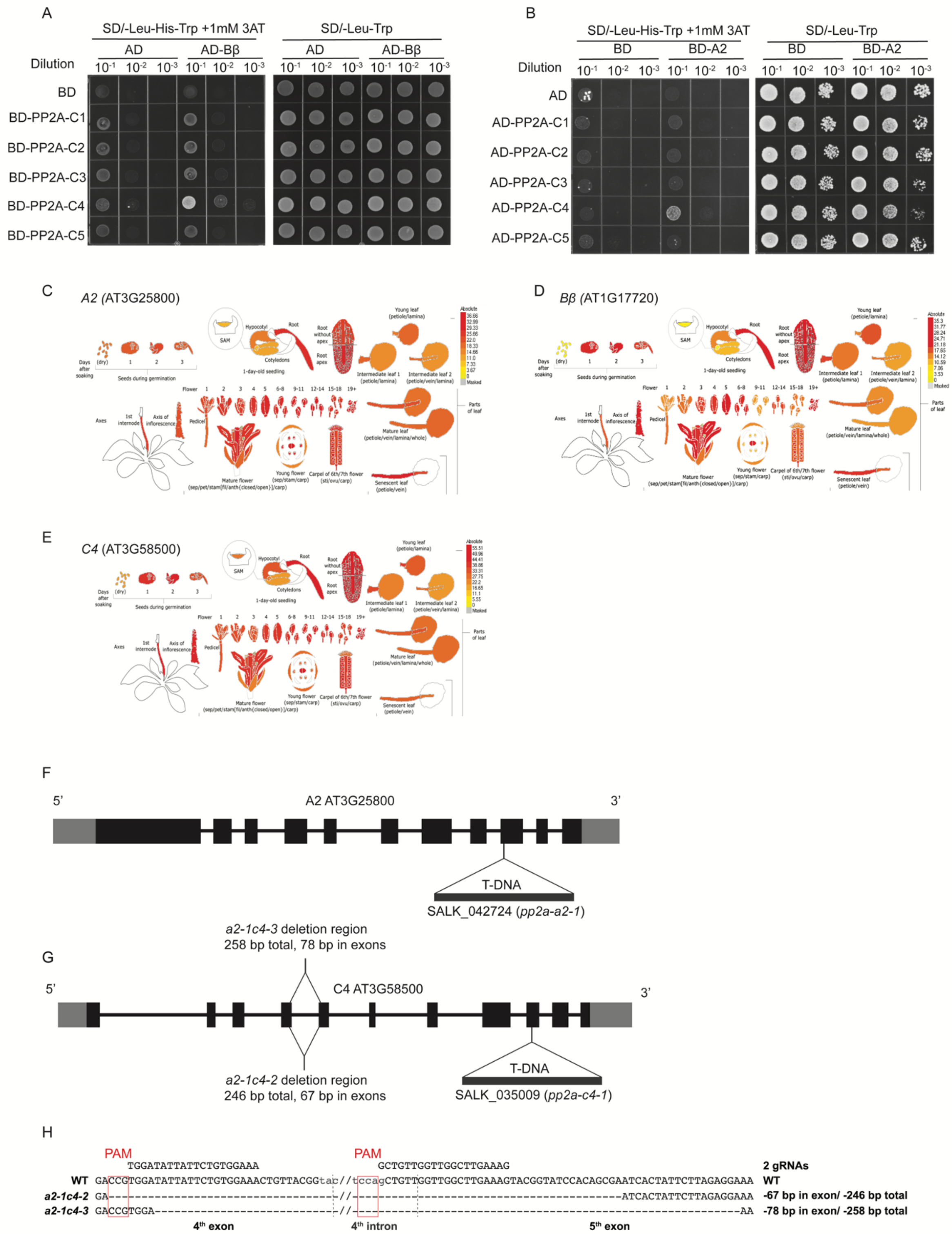


**Figure S2. Related to Figure 2. The interactions between Bβ and C4 or C4 and A2, expression patterns of A2, Bβ, and C4, and PP2A *a2* and *c4* mutants**

(A-B) Yeast two-hybrid assay to screen the interactions of different subunits of PP2A. The interaction between Bβ and C4 (A), and the interaction between A2 and C4 (B). BD: GAL4 DNA binding domain; AD: GAL4 activation domain. Left panel: yeasts grown on selective three-dropout medium; Right panel: yeasts grown on two-dropout medium as a control. (C-E) Expression patterns of A2 (C), Bβ (D), and C4 (E) as illustrated by eFP browser (http://bar.utoronto.ca/efp/cgi-bin/efpWeb.cgi). (F) Schematic diagram to show the genomic structure of *A2* gene (AT3G25800) and the location of the T-DNA insertion in SALK_042724 (*pp2a-a2-1*). (G) Diagram to show the genomic structure of *C4* gene (AT3G58500) and the location of the T-DNA insertion in SALK_042724 (*pp2a-c4-1*), and the deletions in *C4* created by CRISPR-Cas9 in *a2-1* mutant background (*a2-1c4-2, a2-1c4-3*). (H) Locations of the *C4* gRNAs and sequences of mutations in *C4* gene in *a2-1c4-2* and *a2-1c4-3*. Deletions are shown as dashes. The numbers of deletion (base pair) in the exons and in the gDNA for each mutation are shown on the right.

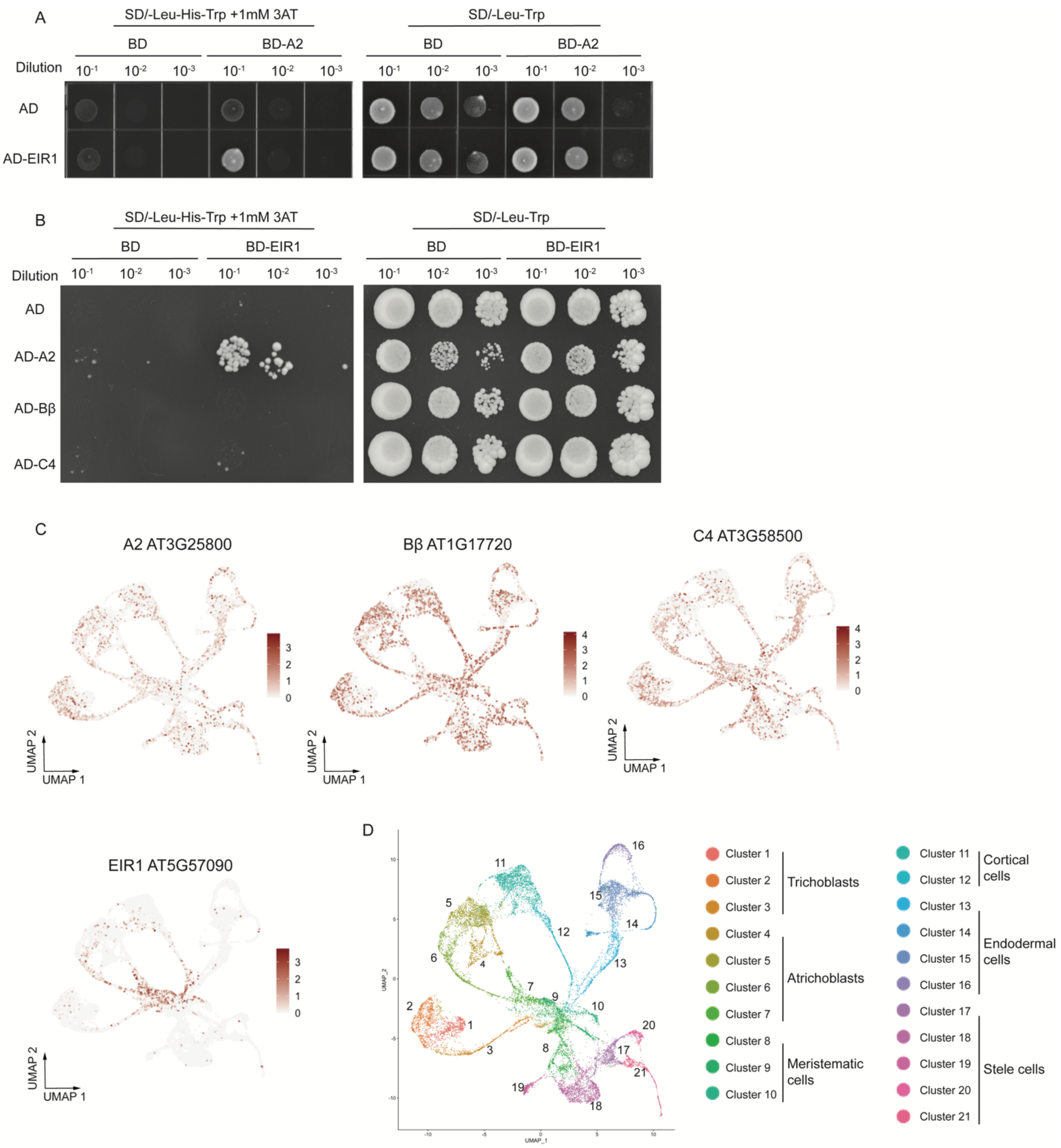


**Figure S3. Related to Figure 3. EIR1 interacts with A2 and the UMAP visualizations of spatial expression profiles for A2, Bβ, C4, and EIR1 in sNucRNA-seq**

(A-B) Yeast two-hybrid assay to test the interaction between EIR1 and A2, C4 or Bβ. The interaction between A2 and EIR1 (A). The interaction between EIR1 and A2, C4 or Bβ (B). BD: GAL4 DNA binding domain; AD: GAL4 activation domain. Left panel: yeasts grown on selective three-dropout medium; Right panel: yeasts grown on two-dropout medium as a control. (C) The expression profile of A2, Bβ, C4, and EIR1 in sNucRNA-seq dataset visualized on the Uniform Manifold Approximation and Projection (UMAP). Aforementioned genes are co-expressed in cluster 3, 6, 7, 9, and 10. (D) UMAP illustration of 21 different *Arabidopsis* root cell clusters from the public sNucRNA-seq dataset with characterized cell types.

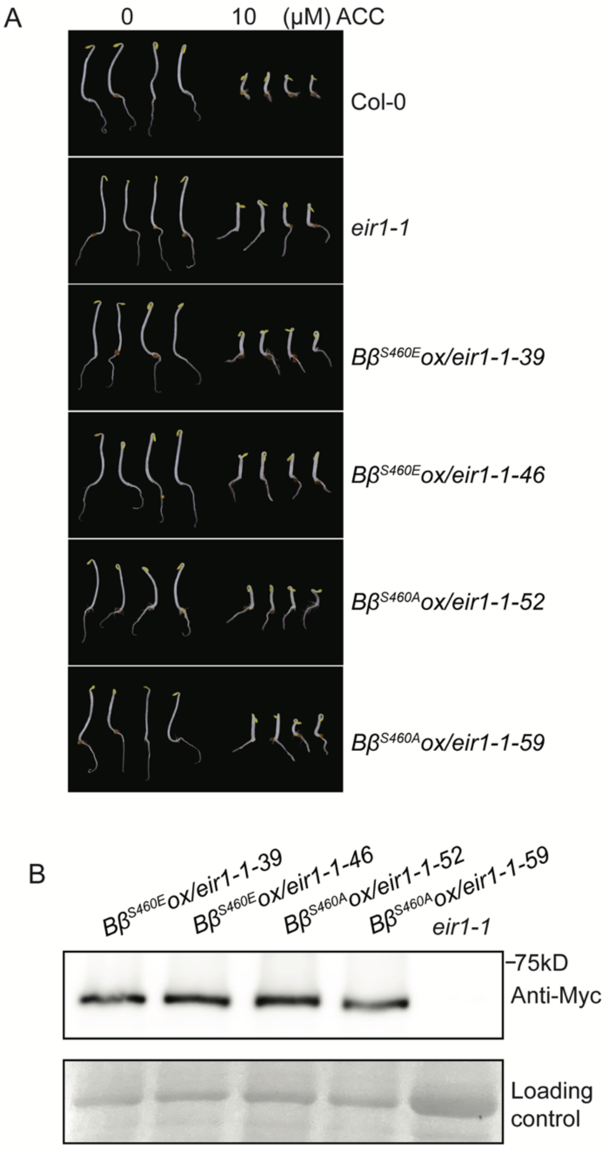


**Figure S4. Related to Figure 5. Bβ regulates root inhibition through EIR1 in response to ethylene**

(A) The ethylene responsive phenotype of the indicated plants. 3-day-old seedlings were grown on MS medium containing 10μM ACC or without ACC in the dark before being photographed. (B) Western blot assay to examine the protein expression in the independent transgenic plants. The total proteins from the indicated transgenic plants were subjected to the blot assay with anti-Myc antibody Ponceau staining serves as a loading control.

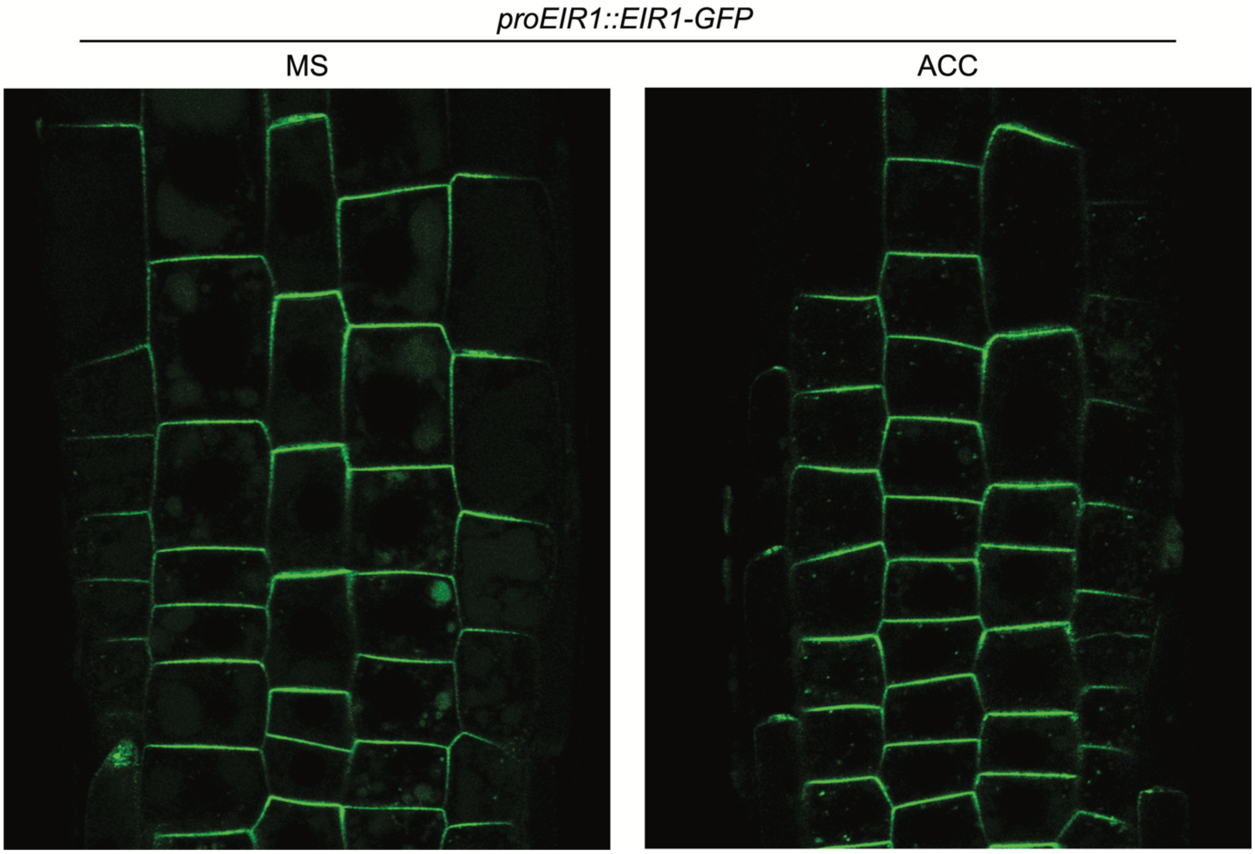


**Figure S5**. **The subcellular localization of EIR1 with or without ethylene treatment.** 3-day-old etiolated seedlings were grown on MS medium with or without 10μM ACC before imagined under confocal microscopy.

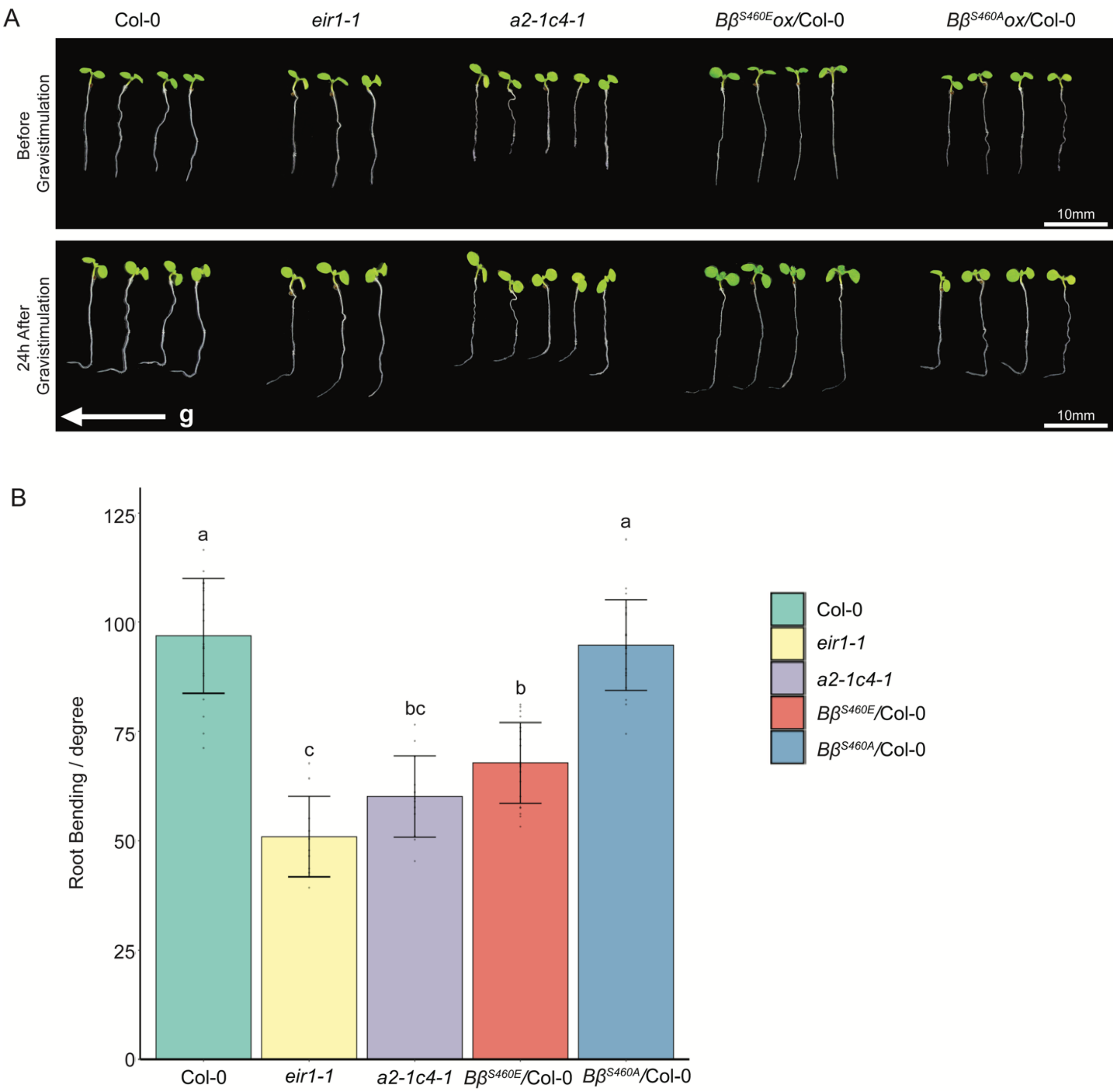


**Figure S6**. **The gravitropism assays of Col-0, *eir1-1*, *a2-1c4-1,* and *Bβ*$^{S460E}$*ox*/Col-0.** (A) Root gravitropic response assay of the indicated plants. Pictures were taken prior to and 24 hours after the gravitropism assays from representative individual plants. Scale bar represents 10mm. (B) Quantification of the root bending angles in each genotype. Individual data are plotted as dots. Different letters indicate significant differences between different genotypes with $P \leqslant 0.05$ calculated by a two-tailed *t* test.
